# Models that learn how humans learn: the case of decision-making and its disorders

**DOI:** 10.1101/285221

**Authors:** Amir Dezfouli, Kristi Griffiths, Fabio Ramos, Peter Dayan, Bernard W. Balleine

## Abstract

Popular computational models of decision-making make specific assumptions about learning processes that may cause them to underfit observed behaviours. Here we suggest an alternative method using recurrent neural networks (RNNs) to generate a flexible family of models that have sufficient capacity to represent the complex learning and decision-making strategies used by humans. In this approach, an RNN is trained to predict the next action that a subject will take in a decision-making task and, in this way, learns to imitate the processes underlying subjects’ choices and their learning abilities. We demonstrate the benefits of this approach using a new dataset drawn from patients with either unipolar (n=34) or bipolar (n=33) depression and matched healthy controls (n=34) making decisions on a two-armed bandit task. The results indicate that this new approach is better than baseline reinforcement-learning methods in terms of overall performance and its capacity to predict subjects’ choices. We show that the model can be interpreted using off-policy simulations and thereby provides a novel clustering of subjects’ learning processes – something that often eludes traditional approaches to modelling and behavioural analysis.

## Introduction

A computational model of decision-making is a mathematical function that inputs past experiences (such as chosen actions and the value of rewards), and outputs predictions about future actions (e.g., Busemeyer and Diederich, 2010; Daw, 2011; Gold and Shadlen, 2007). Typically, experimenters develop such models by specifying a set of structural assumptions along with free parameters that can be adjusted to capture a range of behaviors. However, this approach can only ever capture learning processes that fall within the boundaries of the assumptions embedded in the model structure. If the actual learning and choice processes used by real human subjects differ from those assumptions, e.g., if a single learning-rate parameter is assumed to update the effects of reward and punishment on action values when they are in fact modulated by different learning-rates, then the model will misfit the data (e.g., Piray et al., 2014). To overcome this problem, the process of computational modelling often involves an iterative process, including additional analyses to assess the assumptions about the behavior, and then the amendation of the structural features of the models to reduce residual fitting error, then new analyses, and so forth. The final model is the simplest one that misfits the least. This iterative process has become standard scientific practice for model development in domains such as cognitive science, computational psychiatry, and model-based analyses of neural data (e.g., Busemeyer and Stout, 2002; Dezfouli et al., 2007; Montague et al., 2012; O’Doherty et al., 2007; Miller et al., 2017; Acuña and Schrater, 2010).

Here we consider an alternative approach that involves minimal assumptions about the underlying learning processes used by subjects, derived from a very flexible class of model that essentially learns-to-learn (Hochreiter et al., 2001; Wang et al., 2016; Duan et al., 2016; Weinstein and Botvinick, 2017)^1^. We consider recurrent neural networks (RNNs) as our flexible class, which are known to have sufficient capacity to represent any form of computational process (Siegelmann and Sontag, 1995), including the ones believed to be behind the behaviours of humans and other animals in a wide range of decision-making, cognitive and motor tasks (Song et al., 2017; Zhang et al., 2018; Miconi, 2017; Carnevale et al., 2015; Mante et al., 2013; Song et al., 2016; Barak et al., 2013; Yang et al., 2017; Sussillo et al., 2015; Hennequin et al., 2014; Rajan et al., 2016; Laje and Buonomano, 2013). Learning to learn involves adjusting the weights in a network so that it can predict the choices that subjects make both during learning and at asymptote. After this, the weights are frozen, and the model is simulated on the actual learning task to assess its predictive capacity and to gain insights into the subjects’ behavior. Since these models are flexible, they can automatically characterize the major behavioral trends exhibited by real subjects without requiring tweaking and engineering based on behavioral analyses, something that is particularly useful when major trends in the data are not apparent in behavioral summary statistics. The potential problem is that the models are so flexible that they might not generalize in a relevant manner.

To illustrate and evaluate this approach, we focus on a relatively simple decision-making task, involving a two-arm bandit, in which subjects had a choice between two actions (button presses) that were rewarded probabilistically. To examine the predictive capacity of RNNs under normal and abnormal conditions, data from three groups were collected: healthy subjects, and patients with either unipolar or bipolar depression. We found that RNNs were able to learn the subjects’ decision-making strategies more accurately than baseline reinforcement-learning models. Furthermore, we show that off-policy simulations of the RNN model allowed us to visualize, and thus uncover, the properties of the learning process behind subjects’ actions and that these were inconsistent with the assumptions made by reinforcement-learning treatments. Furthermore, we illustrate how the RNN method can be applied to predict diagnostic categories for different patient populations.

## Results

### Model and task settings

rnn. The architecture we used is depicted in Figure 1; it is a particular form of recurrent neural network. The model is composed of an lstm layer (Long short-term memory; Hochreiter and Schmidhuber, 1997), which is a recurrent neural network, including an output softmax layer with two nodes (since there are two actions in the task). The inputs to the model on each trial are the previous action and the reward received after taking the action, and the outputs of the model are the probabilities of selecting each action in the next trial. We refer to the framework proposed here as rnn.

**Figure 1.**
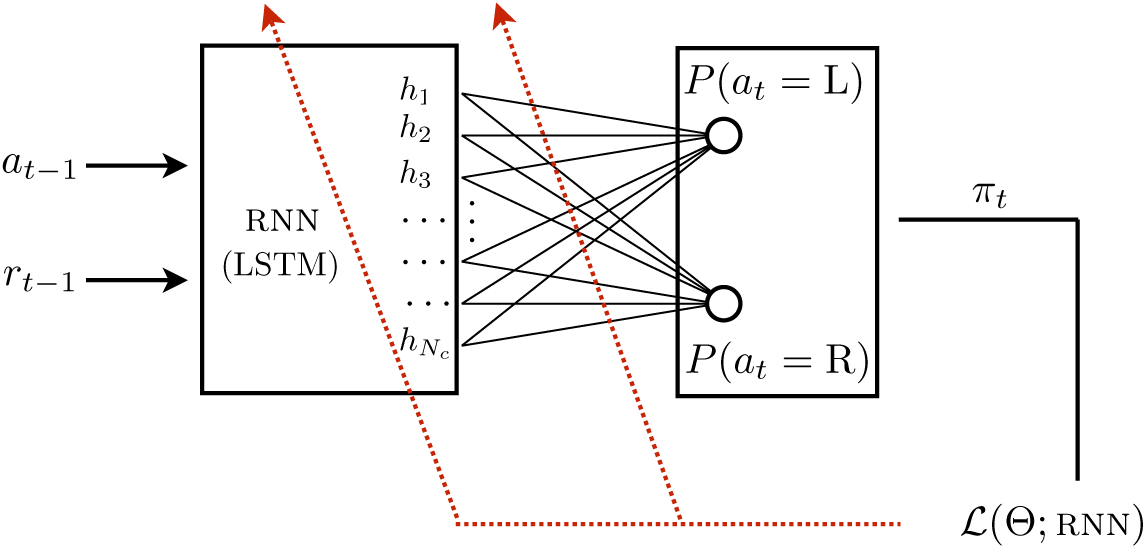
Structure of the rnn model. The model has a lstm layer which receives previous actions and rewards as inputs, and is connected to a softmax layer which outputs the probability of selecting each action on the next trial (policy). The maximum likelihood estimate method was used to train the weights in the lstm and softmax layers. *h_i_* is the output of lstm cell *i* in the lstm layer. *N_c_* is the number of cells in the lstm layer. The dotted red line indicates that information about the metric is used to adjust the weights in the lstm and output layers as the network learns-to-learn.

The lstm layer is composed of a set of interconnected lstm cells, in which each cell can be thought of as a memory unit which maintains and updates a scalar value over time (shown by *h_i_* in Figure 1 for the i*th* lstm cell). On each trial, the value of each cell is updated based on the inputs and on the last value of the other lstm cells in the network (including the cell itself), and in this way the lstm layer can track relevant information regarding the history of past rewards and actions. The nature of the information tracked depends on how lstm cell values are updated, which is modulated by the weights of the connections between the cells and also between the inputs and the cells. As such, the lstm layer is composed of two sets of connections linking lstm cells to each other and also to the inputs. In addition, each lstm cell outputs its current value (*h_i_*) to the softmax layer through a set of connections that determine the influence of the output of each cell on the predictions for the next action (shown by lines connecting them in Figure 1). As a whole, such an architecture is able to *learn* in a decision-making task by tracking the history of past experiences using the lstm layer, and then turning this information into subsequent actions through the outputs of the softmax layer.

The way in which a network learns in the task and maps past experiences to future actions is modulated by weights in the network. Here, our aim was to tune the weights so that the network could predict the next action taken by the subjects – given that the inputs to the network were the same as those that the subjects received on the task. This is *learning-to-learn*, shown by the red arrows in Figure 1, in which the weights are trained to optimise a metric which represents how well the model can predict subjects’ choices (denoted by 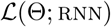 in the figure).

**Baseline models**. We compared the predictive accuracy of the rnn model with classical exemplars from the reinforcement learning (RL) family. The first baseline RL model was the *Q*-learning model (denoted by ql), in which subjects’ choices are determined in a noisy manner by learned action values (often called *Q* values; Watkins, 1989), which are updated based on the experience of reward. The second baseline model was *Q*-learning with perseveration (denoted by qlp), which is similar to ql but has an extra parameter that allows for a tendency to stick with the same action for multiple trials (i.e., to perseverate), or sometimes to alternate between the actions (independent of reward effects; Lau and Glimcher, 2005). As we show below, both ql and qlp offered significantly worse accounts of subjects’ choices than rnn, and so we developed a new baseline model that we called generalised *Q*-learning (denoted by gql). This model extends ql and qlp models by learning multiple values for each action using different learning rates, and also by tracking the history of past actions at different time scales.

**Task and subjects**. The instrumental learning task (Figure 2) involved participants choosing between pressing a left (L) or right (R) button (self-paced responses) in order to earn food rewards (an M&M chocolate or a BBQ flavoured cracker). During each 40 second block, one action was always associated with a higher probability of reward than the other (which always had the value 0.05). Across 12 blocks, the action with the higher reward probability switched identities (left or right), and the probability was 0.25, 0.125, or 0.08, drawn at random. 34 uni-polar depression (depression), 33 bipolar (bipolar) and 34 control (healthy) participants (age, gender, IQ and education matched) completed the task. See Materials and methods for the details of the task and the models.

**Figure 2.**
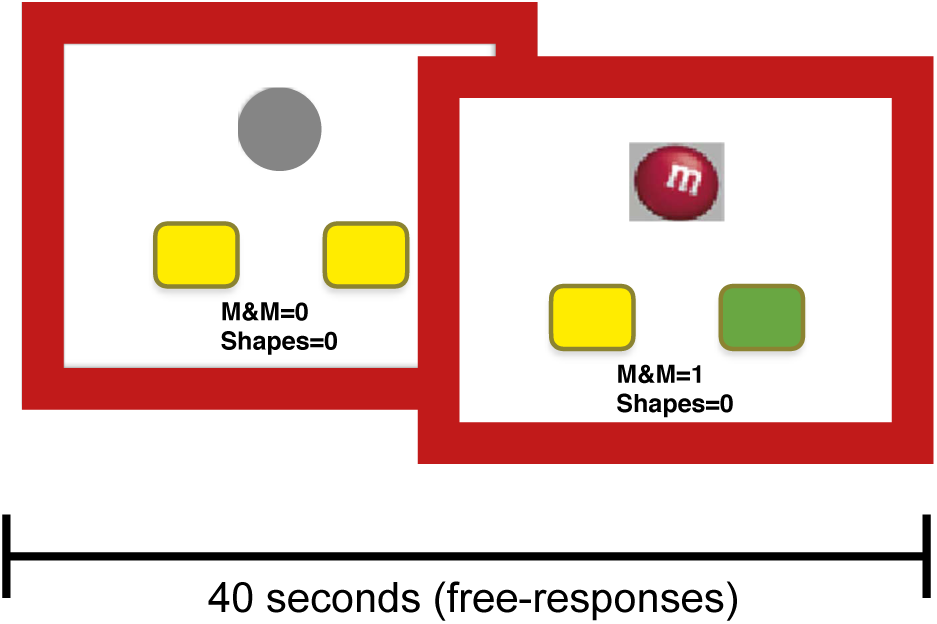
Structure of the decision-making task. Before the choice, no indication was given as to which button was more likely to lead to reward. When the participant made a rewarded choice, the button chosen was highlighted (green) and a picture of the earned reward was presented for 500ms. Each block lasted for 40 seconds and each participant completed 12 blocks. Blocks were separated by a 12-second inter-block interval.

### Performance in the task

We start by describing the high level properties of subjects’ choices. Figure 3 shows the probability of selecting the best action (i.e., the action with the higher reward probability). Results for subjects are shown by subj in the graph. The probability of selecting the better action was significantly higher than the other action in all groups (healthy [*β* = 0.270, SE=0.026, *p<* 0.001]^2^, depression [*β* = 0.149, SE=0.028, *p<* 0.001], bipolar [*β* = 0.119, SE=0.021, *p<* 0.001]). Comparing healthy and depression groups, revealed that the group by action interaction had a significant effect on the probability of selecting actions [*β* = −0.120, SE=0.038, *p* = 0.002]^3^. A similar effect was observed when comparing the healthy and bipolar groups [*β* = −0.150, SE=0.034, *p<* 0.001]. In summary, these results indicate that all groups were able to direct their actions toward the better choice, however the depression and bipolar groups were less able to do so compared to the healthy group.

**Figure 3.**
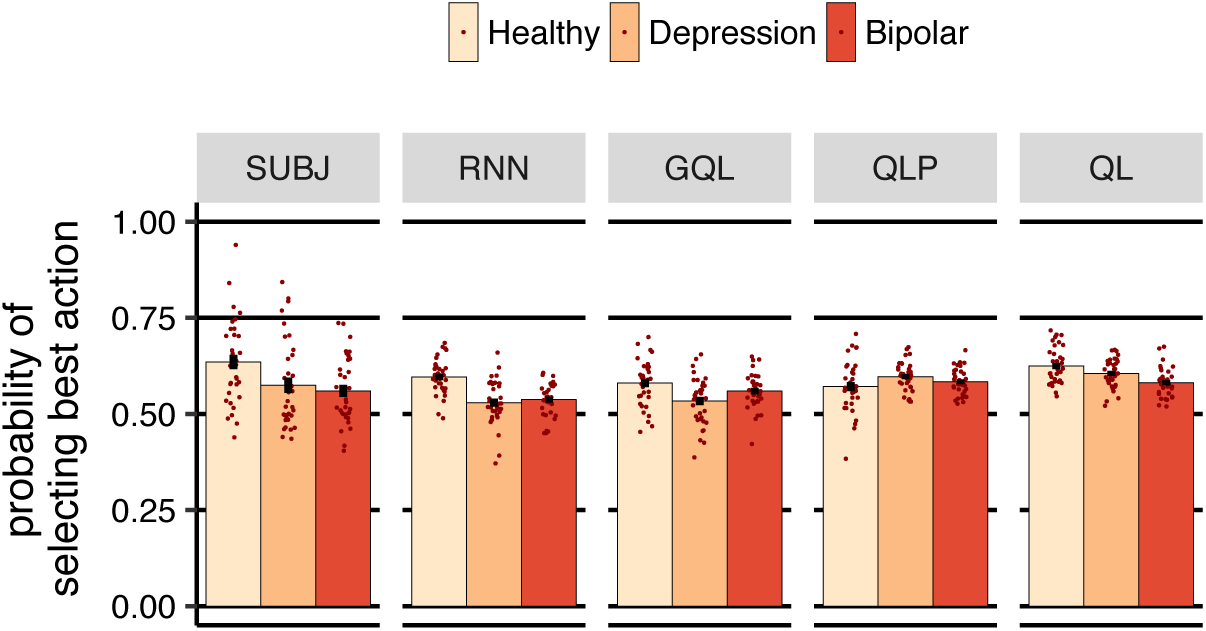
Probability of selecting the action with the higher reward probability (averaged over subjects). Each dot represents a subject and error-bars represent 1 SEM.

Next, we trained three instances of rnn using the data from each group, and then froze the weights of the models and simulated them on-policy in the task (with the same reward probabilities and for the same number of trials that each subject completed). On-policy means that the models completed the task on their own by selecting the actions that they predicted a representative subject would take in each situation. The results of the simulations are shown in Figure 3 in the rnn column. Similar to the subjects’ data, the probability of selecting the better action was significantly higher than the other action in all the three groups (healthy [*β* = 0.192, SE=0.011, *p<* 0.001], depression [*β* = 0.058, SE=0.014, *p<* 0.001], bipolar [*β* = 0.074, SE=0.011, *p<* 0.001]). Therefore, although the structure of rnn was initially unaware that the objective of the task was to collect rewards, its actions were directed toward the better key by following the strategy that it learned from the subjects’ actions. A similar pattern was observed for gql, qlp and ql models in the figure, which is not surprising as the structure of these models includes value representations which can be used for reward maximization (estimated parameters for ql, qlp and gql models are shown in Tables S2, S3 and S4 respectively; the negative log-likelihood for each model is reported in Table S5. See Table S7 for the effect of the initialisation of the network on the negative log-likelihood of trained rnn. See Table S6 for the negative log-likelihood when a separate model was fitted to each subject in the case of baseline models).

### The immediate effect of reward on choice

Based on the analyses described in the previous section, similarly to the subjects, rnn was able to guide its actions toward the better choices. However, there are multiple strategies that the models could follow to achieve this, and here we aimed to establish whether the strategy used by the models was similar to that used by the subjects’. We started by investigating the immediate effect of reward on choices. Figure 4 shows the effect of earning a reward on the previous trial on the probability of staying on the same action in the next trial. For the subjects (subj), earning a reward significantly *decreased* the probability of staying on the same action in the healthy and depression groups, but not in the bipolar group (healthy [*β* = 0.112, SE=0.019, *p<* 0.001]^4^, depression [*β* = 0.111, SE=0.029, *p<* 0.001], bipolar [*β* = 0.030, SE=0.035, *p* = 0.391]). As the figure shows, the same pattern was observed in rnn (healthy [*β* = 0.082, SE=0.006, *p<* 0.001], depression [*β* = 0.089, SE=0.013, *p<* 0.001], bipolar [*β* = 0.001, SE=0.010, *p* = 0.887]), which shows that the strategy used by rnn was similar to the subjects’ according to this analysis.

**Figure 4.**
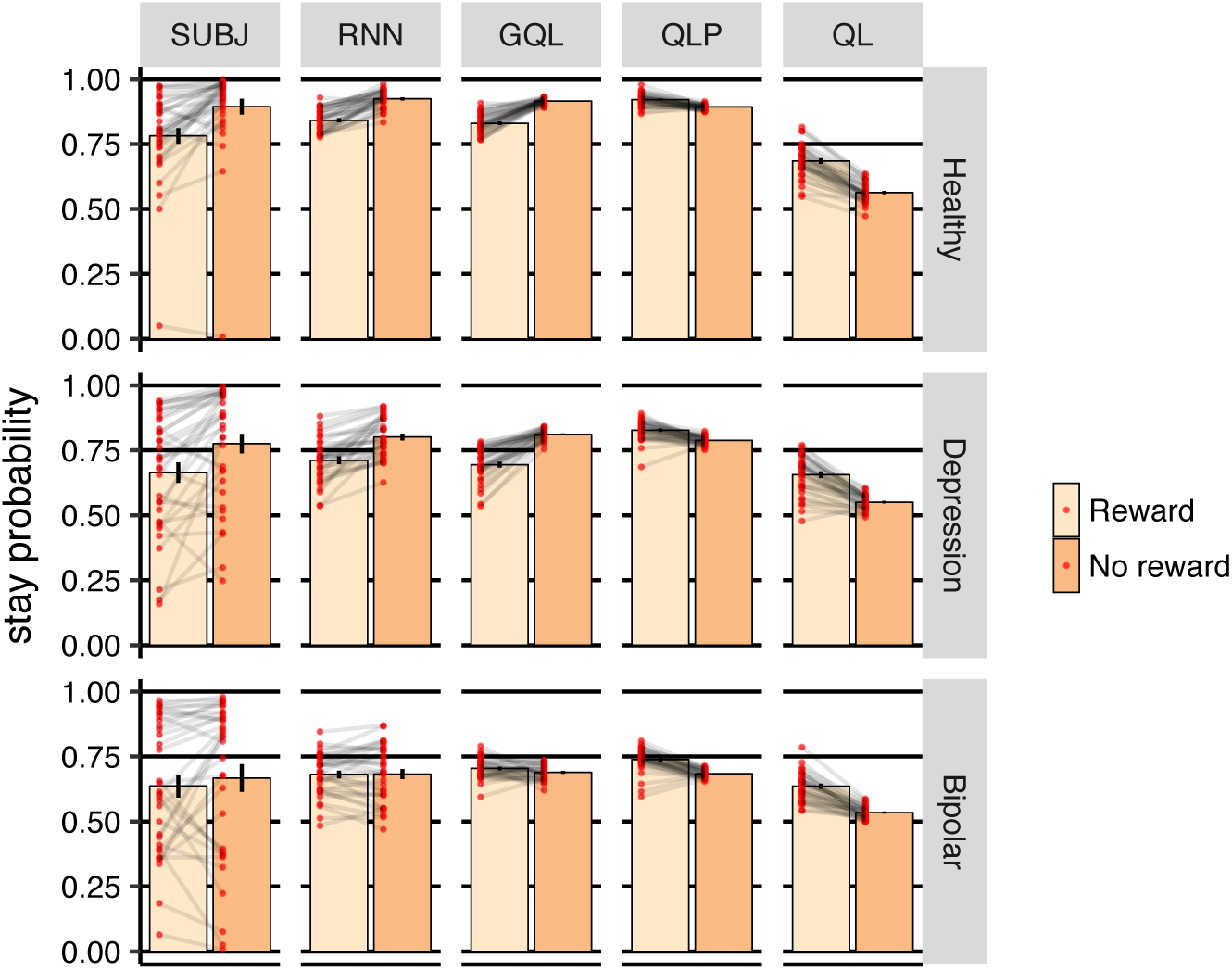
Probability of staying on the same action based on whether the previous trial was rewarded (Reward) or not rewarded (No reward), averaged over subjects. Each dot represents a subject and error-bars represent 1SEM.

In contrast, stay probabilities had opposite directions in ql and qlp, i.e., the probability of staying on the same action was *higher* after earning reward (for the case of qlp; healthy [*β* = −0.028, SE=0.004, *p<* 0.001], depression [*β* = −0.039, SE=0.006, *p<* 0.001], bipolar [*β* = −0.054, SE=0.007, *p<* 0.001]), which differs from the subjects’ data. This pattern was expected from baseline reinforcement-learning models, i.e., ql and qlp as, in these models, earning reward increases the value of the taken action, which raises the probability of choosing that action on the next trial. Indeed, this learning process is embedded in the parametric forms of ql and qlp models, and cannot be reversed no matter what values are assigned to the free-parameters of these models. As such, we designed gql as a baseline model with more relaxed assumptions, in which action values could have an opposite effect on the probability of selecting actions, and so could generate a similar of responding pattern to the subjects’.

Despite the fact that gql could capture the high-level behavioural summaries, it remains an open question whether this model could represent all the behavioural trends in the data, or whether there were some missing trends undetectable in the summary statistics presented here. In the next section we will answer this question by comparing the prediction capacity of gql with rnn, as a model that has the potential capacity to capture all of the behavioural trends in the data. A second question relates to the strategy that the subjects are using. It is not immediately clear how choice was directed toward the better action while, at the same time, the probability of switching to the other action after earning rewards was higher, some that implies that choice was diverted from the better action. We answer this latter question using off-policy simulations of the models in the following sections.

### Action prediction

Here our aim was to quantify how well the models predicted the actions chosen by the subjects. We used Leave-one-out cross-validation for this purpose in which, at each round, one of the subjects was withheld and the model was trained using the remaining subjects; the trained model was then used to make predictions about the withheld subject. The withheld subject was rotated in each group, yielding 34, 34 and 33 prediction accuracy measures in the healthy, depression, and bipolar groups, respectively.

The results are reported in Figure 5. The left-panel of the figure shows prediction accuracy in terms of nlp (negative log-probability; averaged over leave one-out cross-validation folds; lower values are better) and the right-panel shows the percentage of actions predicted correctly (‘%correct’; higher values are better). nlp roughly represents how well each model fits the choices of the withheld subject, and unlike ‘%correct’, it takes the certainty of predictions into account. Therefore, we focus on nlp in the analysis. Firstly, gql had the best nlp among the baseline models in all the three groups; its advantage was statistically significant in the depression and bipolar groups (healthy [*β* = −0.036, SE=0.020, *p* = 0.086]^5^, depression [*β* = −0.101, SE=0.024, *p<* 0.001], bipolar [*β* = −0.105, SE=0.019, *p<* 0.001]). Secondly, rnn’s nlp was even better than gql across all groups (healthy [*β* = 0.090, SE=0.040, *p* = 0.030], depression [*β* = 0.126, SE=0.021, *p<* 0.001], bipolar [*β* = 0.180, SE=0.032, *p<* 0.001], showing that rnn was able to predict subjects’ choices better than the other models.

**Figure 5.**
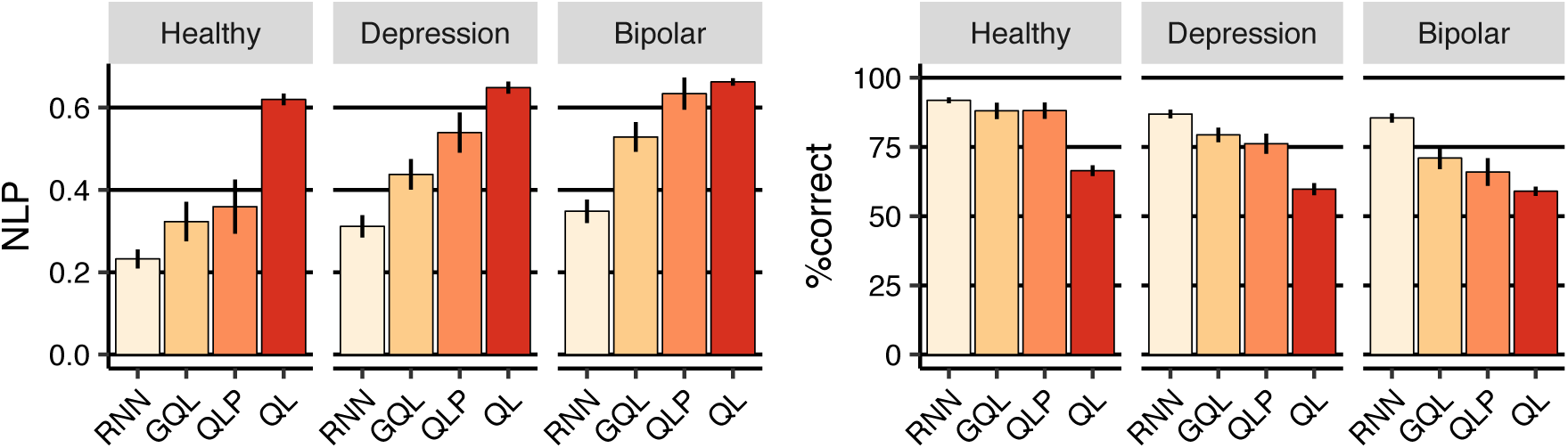
Cross-validation results. **(Left-panel)** nlp (negative log-probability) averaged across leave-one-out cross-validation folds. Lower values are better. **(Right-panel)** Percentage of actions predicted correctly averaged over cross-validation folds. Error-bars represent 1SEM.

The fact that gql is better than ql and qlp is not unexpected; we showed in the previous section that the predictions from ql and qlp were inconsistent with the trial-by-trial behaviour of the subjects. On the other hand, the fact that rnn is better than gql shows that there are some behavioural trends that even gql failed to capture, although it was consistent with subjects’ choices according to the behavioural summary statistics. In the next sections, we will use off-policy simulations of the models to uncover the additional behavioural trends that were captured by rnn.

### Off-policy simulations

In an off-policy simulation, a model uses information about previous choices and rewards to make predictions about the next action; but the actual next action is not derived from its predictions, but are rather determined in some other manner (notably, from human choices). In this way we can control what inputs the model receives and examine how they affect predictions. We were interested, in particular, in establishing how the predictions of the models were affected by the history of previous actions and rewards. As such, we designed a variety of inputs based on the behavioural statistics, fed them into the models, and recorded the predictions of the model in response to each input set (see Section S2 for more details on how simulation parameters were chosen). Simulations of the models (rows) are shown in Figure 6 for the healthy group, in which each panel shows a separate simulation across 30 trials (horizontal axis). For trials 1–10, the action that was fed to the model was R, and for trials 11–30 it was L (the action fed into the model at each trial is shown in the ribbons below each panel). The rewards associated with these trials varied among simulations (the columns) and are shown by black crosses (x) in the graphs.

**Figure 6.**
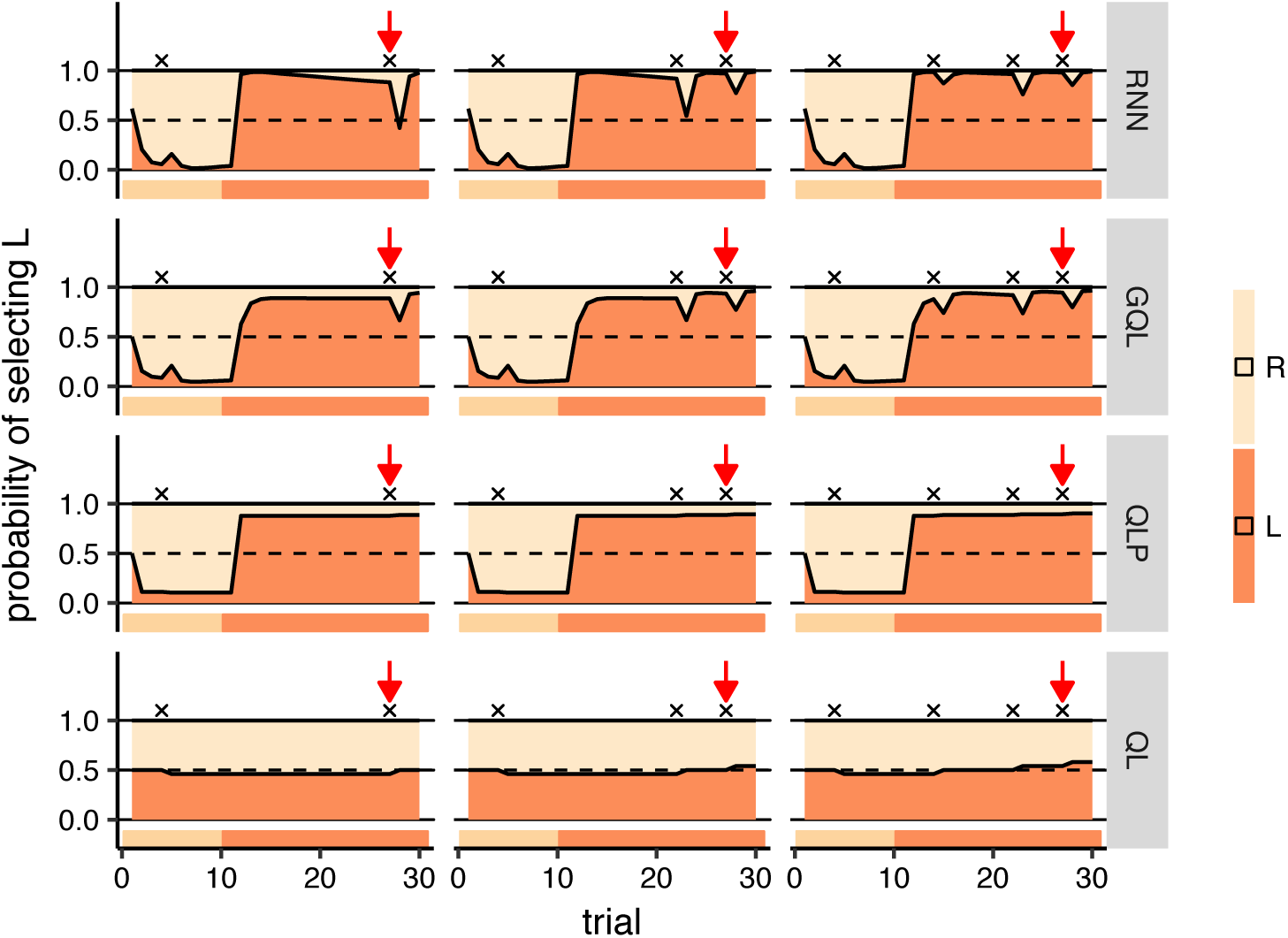
Off-policy simulations of all models for group healthy. Each panel shows a simulation for 30 trials (horizontal axis), and the vertical axis shows the predictions of each model on each trial. The ribbon below each panel shows the action which was fed to the model on each trial. In the first 10 trials, the action that the model received was R and in the next 20 trials it was L. Rewards are shown by black crosses (x) on the graphs. See text for the interpretation of the graph. Note that the models’ prediction for each trial is made *before* seeing which action and reward was fed to the model on that trial.

### The effect of reward on choice

Focusing on the rnn simulations in Figure 6, it can be observed that earning a reward (shown by black crosses) caused a ‘dip’ in the probability of staying with an action, which showed a tendency to switch to the other action. This is consistent with the observation made in Figure 4 that the probability of switching increases after reward. We saw a similar pattern in gql, but in the ql and qlp models the pattern was reversed, i.e., the probability of choosing an action increased after a reward due to an increase in action values (the effects are rather small for qlp and may not be clear for this model), which is again consistent with the observations in Figure 4. The reason that gql was able to produce different predictions to ql and qlp is that, in this model, the contribution of action values to choices can be negative, i.e., higher values can lead to a lower probability of staying with an action (see Section S1 for more explanation).

The next observation was the effect of previous reward on the probability of switching after a reward. First we focused on the rnn model and on the trials shown by red arrows in Figure 6. The red arrows point to the same trial number, but the number of rewards earned prior to the trial differed. As the figure shows, the probability of switching after reward was lower in the right-panel compared to the left and middle panels. The only difference between simulations is that, in the right panel, two more rewards were earned before the red arrow. Therefore, the figure shows that although the probability of switching was higher after reward, it got smaller as more rewards had previously been earned by an action. Indeed, this strategy made subjects switch more from the inferior action as rewards were sparse on that action, and switch less from the superior action, as it was more frequently rewarded. This can reconcile the observations made in Figures 4, 3 that more responses were made on the better action while, at the same time, the probability of switching after reward was higher. Figure 7 shows the same simulations using rnn for all groups. Comparing the predictions at the red arrows for the depression and bipolar groups, we saw a pattern similar to the healthy group, although the differences were smaller in the bipolar group (see Figure S9 for the effect of the initialisation of the model).

**Figure 7.**
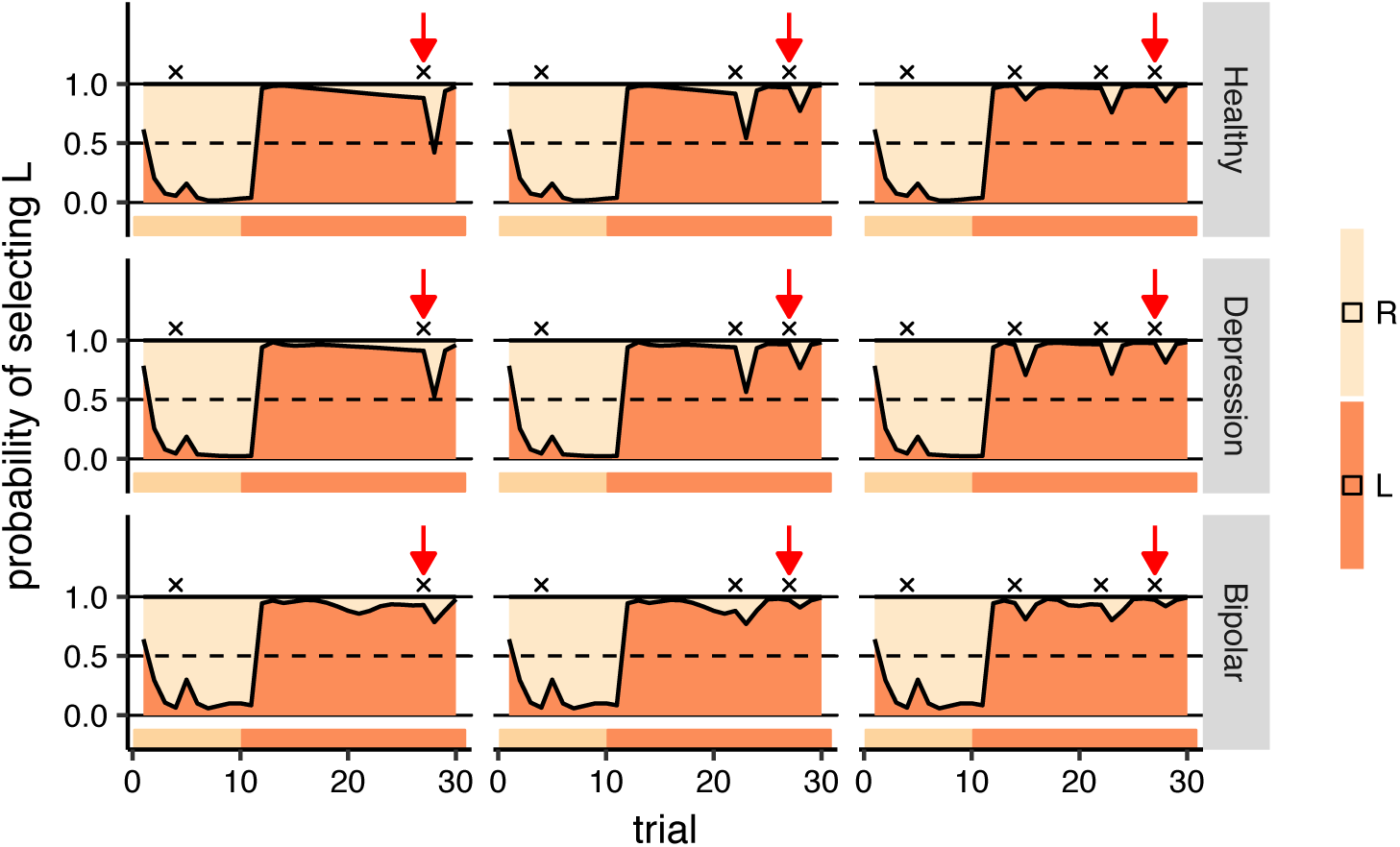
Off-policy simulations of rnn for all groups. Each panel shows a simulation of 30 trials (horizontal axis), and the vertical axis shows the predictions of each model on each trial. The ribbon below each panel shows the action which was fed to the model on each trial. In the first 10 trials, the action that the model received was R and in the next 20 trials it was L. Rewards are shown by black crosses (x) on the graphs. See text for the interpretation of the graph. Note that the simulation conditions are the same as those shown in Figure 6, and the first row here (healthy group) is the same as the first row shown in Figure 6 which is shown here again for the purpose of comparison with the other groups.

The above observations are consistent with the pattern of choices in the empirical data shown in Figure 8-left panel, which depicts the probability of staying with an action after earning reward as a function of how many rewards were earned after switching to the action (a similar graph using on-policy simulation of rnn is shown in Figure S11). In all three groups, the probability of staying with an action (after earning a reward) was significantly higher when more than two rewards were earned previously (>2) compared to when no reward was earned (healthy [*β* = 0.148, SE=0.037, *p<* 0.001]^6^, depression [*β* = 0.188, SE=0.045, *p<* 0.001], bipolar [*β* = 0.150, SE=0.056, *p* = 0.012]), which is consistent with the behaviour of the rnn.

**Figure 8.**
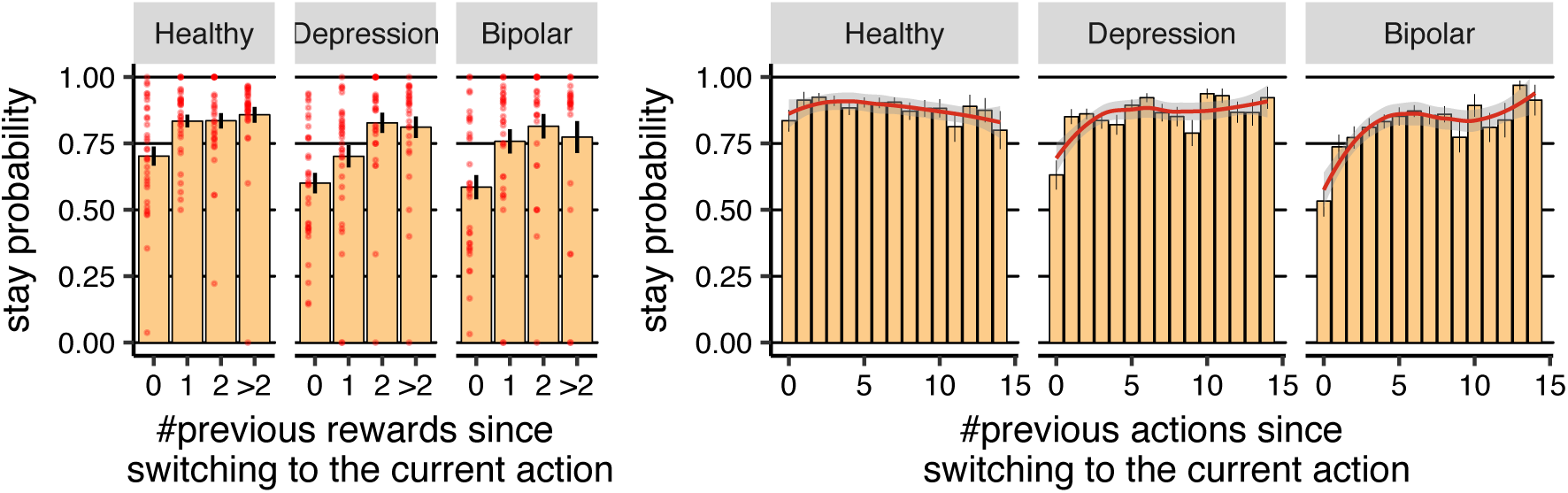
The effect of the history of previous rewards and actions on the future choices of the subjects. **(Left-panel)** The probability of staying with an action after earning reward as a function of the number of actions taken since switching to the current action (averaged over subjects). Each red dot represents the data for each subject. **(Right-panel)** The probability of staying with an action as a function of the number of actions taken since switching to the current action. The red line was obtained using Loess regression (Local Regression), which is a non-parametric regression approach. The grey area around the red line represents the 95% confidence interval. Error-bars represent 1SEM.

As shown in Figure 6, the gql model produced a pattern similar to rnn, which is because this model tracks multiple values for each action, which allows this model to produce the ‘dip’s after rewards, with a magnitude that is sensitive to the number of past rewards (see Section S1 for details. See Figures S5, S6, S7 for gql, qlp and ql models respectively). As such, it is not surprising that gql was consistent with the subjects’ choices with respect to the effect of immediate reward.

**The effect of repeating an action on choices**. Next, we look at the effect of history of actions on choices. Focusing in figure 6 on the rnn model, we can see that, in the first 10 trials, the predicted probability of taking R was higher than L; this then reversed in the next 20 trials. This implies that perseveration (i.e., sticking with the previously taken action) was an element of action selection. This is consistent with the fact that the qlp model (which has a parameter for perseveration) performed better than the ql model in the cross-validation statistics (see Figure 5); and indeed, Figure 6 shows ql’s inability to reflect this characteristic^7^.

Focusing on rnn simulations in the left-panel of Figure 6, we observed that, after switching to action L (after trial 10), the probability of staying with that action gradually decreased, i.e., although there was a high chance the next action would be similar to the previous action, subjects developed a tendency to switch the longer they stayed with an action. To compare this pattern with the empirical data, we calculated the probability of staying with an action as a function of how many times the action had been taken since switching (Figure 8:right-panel)^8^(similar graphs for rnn and gql on-policy simulations are shown in Figures S11, S12 respectively). As the figure shows, for the healthy group, the chance of staying with an action decreased as the action was repeated [*β* = −0.005, SE=0.001, *p* = 0.006]^9^, which is consistent with the behaviour of rnn. With regard to the baseline models, going back to Figure 6, we did not see a similar pattern, although in gql there was a small decrement in the probability of staying with an action after earning the first reward.

**Symmetric oscillations between actions**. Next, we focus on the rnn simulations in Figure 7 in depression and bipolar groups for which the gap between prediction accuracy of gql and rnn was largest. As we see in the left-panels, after switching to action L (after trial 10), the probability of staying with that action gradually decreased in the depression group, but for the bipolar group, there was a dip around 10 trials after switching to action L (i.e., around trial 20), and then the policy became flat. With reference to the empirical data, as shown in Figure 8:right-panel, for the depression and bipolar groups, the probability of staying with an action immediately after switching to that action was around 50% - 60% (shown by the bar at *x* = 0 in Figure 8:right-panel), i.e., there was a 40% - 50% chance that the subject immediately switched back to the previous action. Based on this, we expected to see a ‘dip’ in the simulations of the depression and bipolar groups in Figure 7 just after the switch to action L. This was not the case, pointing to an apparent inconsistency between model predictions and the empirical data.

However, Figure 7 is based on particular, artificial sequences of actions and rewards. To look closer at the above effect, we defined a *run* of actions as a sequence of presses on a certain button without switching to the other button^10^. Figure 9 shows the relationship between consecutive run length, i.e., the length of the current run of actions, as a function of the length of the previous run of actions (see Figures S13, S14, and S15 for similar graphs using on-policy simulations of rnn, gql with *N* = 2 and gql with *N* = 10, respectively). The dashed line in the figure indicates the points at which the current run length was the same as the previous run length. Being close to this line implies that subjects were performing symmetrical oscillations between the two actions, i.e., going back and forth between the two actions while performing an equal number of presses on each button. In particular, as the graph shows in the bipolar group, and to an extent in the depression group, a short run triggered a subsequent run of a similar brevity (see Figures S2, S3, and S4 for raw empirical data). This implies that if, for example by chance, a subject performed a run of length 1, that would initiate a sequence of oscillations between the two actions, keeping the stay probabilities low during short runs, consistent with what was seen at *x* = 0 in Figure 8:right-panel. This effect was not seen in the simulations shown in Figure 7, because the length of the previous run before switching to action L was 10 (there were 10 R actions), and therefore we should not expect the next run to be of length 1, nor should we have actually expected to see a dip in policy just after the first switch.

**Figure 9.**
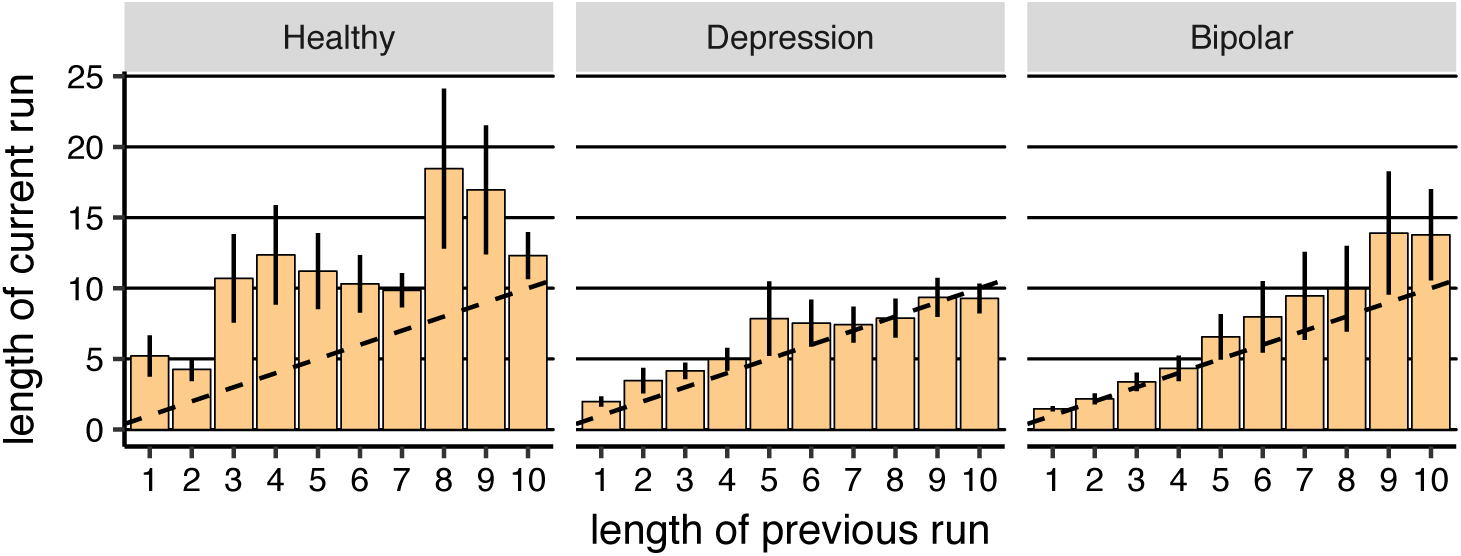
The median number of actions executed sequentially before switching to another action (run of actions) in each subject as a function of the length of the previous run of actions (averaged over subjects). The dotted line shows the points at which the length of the previous and the current run of actions were the same. Note that the median was used instead of the average to illustrate the most common ‘current run length’, instead of the average run length for each subject. Error-bars represent 1SEM.

As shown in Figure S10, the modal lengths of runs in the depression, and bipolar groups were 1 (around 17%, 37%, and 45% of runs were of length 1 in the healthy, depression, and bipolar groups respectively). Given this, and the specific pattern of oscillations in the depression and bipolar groups, our next question was whether, in the models, a run of length 1 triggered oscillations similar to those observed in the empirical data. We used a combination of off-policy and on-policy model simulations to answer this question; i.e., during the off-policy phase we forced the model to make an oscillation between the two actions, and then allowed the model to select between actions. We expected, in the healthy group, that the model would converge on one action, whereas, in the depression and bipolar groups, we expected the initial oscillations to trigger further switches. Simulations are presented in Figure 10, which shows that the sequence of actions fed to the model for the first 9 (off-policy) trials was:

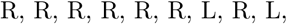
in which there were two oscillations at the tail of the sequence (R, L, R, L,). The rest of the actions (trials 10–20) were selected based on which action the model assigned the highest probability^11^. As the simulation shows, at the beginning, the probability the model assigned to action R was high, but after feeding in the oscillations, the model predicted that the future actions would oscillate in the depression and bipolar groups, but not in the healthy group, consistent with what we expected to observe.

**Figure 10.**
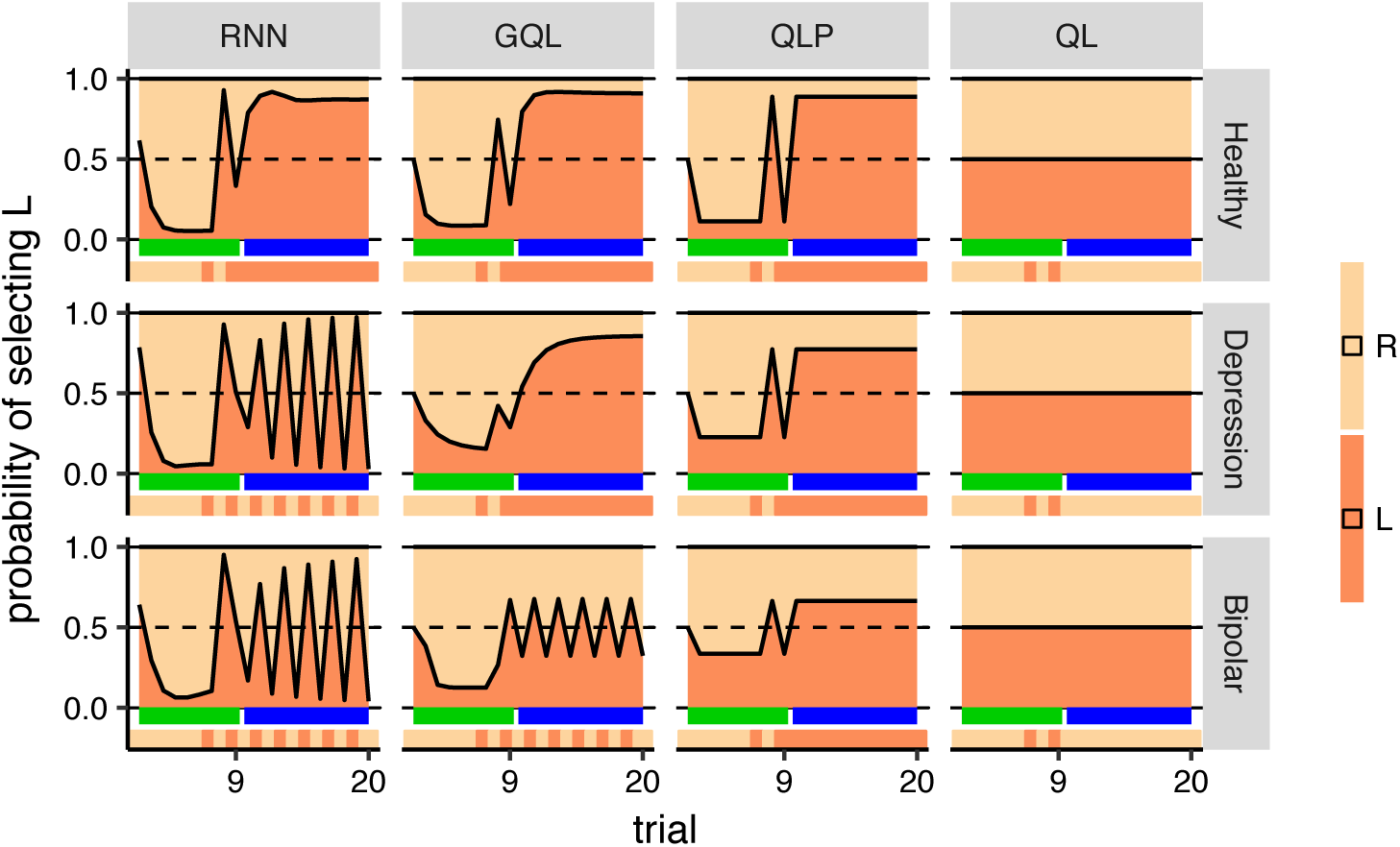
Mixed off-policy and on-policy simulations of the models. Each panel shows a simulation of 20 trials, for which the first nine trials were off-policy and the next trials were on-policy, during which the action with the highest probability was selected. Trials marked with the green ribbons were off-policy (actions were fed to the model), whereas the trials marked with the blue ribbons were on-policy (actions are selected by the model). The ribbon below each panel shows the actions that were fed to the model (for the first 9 trials), and the actions that were selected by the model (for the rest of trials). During off-policy trials, the sequence of actions that was fed to the model was R, R, R, R, R, R, L, R, L. See the text for interpretation.

Therefore, rnn was able to produce symmetrical oscillations and its behaviour was consistent with the subjects’ actions. As Figure 10 shows, besides rnn, gql was also able to produce length 1 oscillations to some extent (as shown for the bipolar group), which could explain why the prediction accuracy achieved by this model was significantly better than qlp in the bipolar and depression groups (Figure 5) in which length 1 oscillations were more common (see Section S2 for more details). However, as shown in Figure S14, gql failed to produce oscillations of longer lengths (even if we increased the capacity of gql to track 10 different values for each action; see Figure S15), whereas rnn was able to do so (Figure S13). This inability of the gql model is particularly problematic in the depression and healthy groups, because these two groups tended to match the length of consecutive runs of actions. This could partly account for why the cross-validation statistics associated with rnn were significantly better than gql for the depression and bipolar groups.

**Summary**. Firstly, we found that a rnn model was able to capture the immediate effect of rewards on actions (i.e., the ‘dip’ after rewards), as well as the effect of previous rewards on choices. gql had a similar ability, which enabled it to reproduce the behavioural summary statistics shown in Figures 3, 4. Baseline reinforcement-learning models (qlp and ql) failed to capture either trend. Secondly, rnn was able to capture how choices change as an action is chosen repeatedly and sequentially, and also the symmetrical oscillations between actions, neither of which could be detected by gql.

### Diagnostic label prediction

In the previous sections we showed that there are several behavioural trends that baseline models failed to capture. Here we asked whether capturing such behavioural trends in this task is necessary to predict the diagnostic labels of the subjects. We used the leave-one-out cross-validation method in which, in each run, one of the subjects in each group was withheld, and a rnn model was fitted to the rest of the group. This model, along with the versions of the same model fitted to all the subjects in each of the other two groups, was used to predict the diagnostic label for the withheld subject. This prediction was based on which of the three models provided the best fit (lowest nlp) for that subject. The results are reported in Table 1. Baseline random performance was near 33%. As the table shows, the highest performance was achieved for the healthy group of which 64% of subjects were classified correctly. On the other hand, in the depression group a significant portion of subjects were classified as healthy. The overall correct classification rate of the model was 52%, whereas gql achieved 50% accuracy (Table S1). We conclude, therefore, that although gql was unable accurately to characterize behavioural trends in the data, the group differences that were captured by gql appeared sufficient to guide diagnostic label predictions.

## Discussion

We used a recurrent neural network to provide a framework for learning a computational model that can characterize human learning processes in decision-making tasks. Unlike previous work, the current approach makes minimal assumptions about these learning processes; we showed that this agnosticism is important in developing an appropriate explanation of the data. In particular, the rnn model was able to encode the melange of processes that subjects appeared to use to select actions; it was also able to capture differences between the psychiatric groups. These processes were largely inconsistent with conventional and tailored *Q*-learning models, and were also hidden in the overall performance of subjects on the task. This provided a clear example of how the currently proposed framework can outperform previous approaches.

**Table 1.**
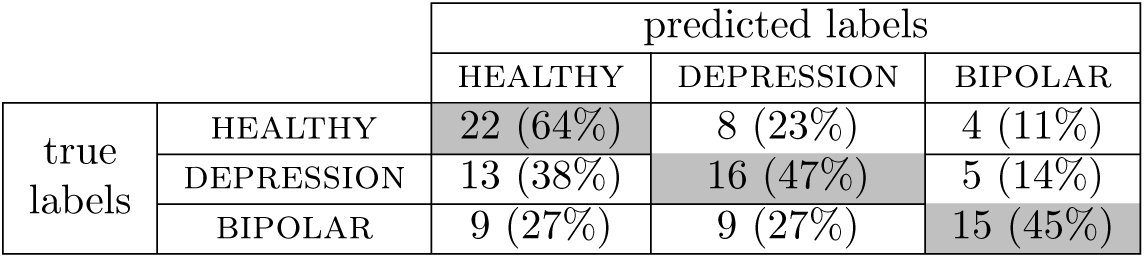
Prediction of diagnostic labels using rnn. Number of subjects for each true-and predicted-labels. The numbers inside the parentheses are the percentage of subjects relative to the total number of subjects in each diagnostic group.

|  |  | predicted labels |  |  |
| --- | --- | --- | --- | --- |
|  |  | HEALTHY | DEPRESSION | BIPOLAR |
| true labels | HEALTHY | 22 (64%) | 8 (23%) | 4 (11%) |
|  | DEPRESSION | 13 (38%) | 16 (47%) | 5 (14%) |
|  | BIPOLAR | 9 (27%) | 9 (27%) | 15 (45%) |

In general, as we were able to show, this new approach improves upon previous methods from four standpoints: First, it provides a model that was able to predict subjects’ choices without requiring manual engineering; and to do so more accurately than baseline models on validation data. Second, the framework contributes to computational modelling by providing a baseline for predictive accuracy; i.e., to the extent that other candidate models failed to generate the performance of rnn models, important and accessible behavioural trends would have been missed in the model structure. This is particularly important because of the natural randomness of human choice, making it unclear in many scenarios whether the model at hand (e.g., a *Q*-learning model) has reached a limit as to how well those choices can be predicted, or whether it requires further improvements. Without other recourse, conventional treatments tend to relegate model mis-fit to irreducible randomness in choice. Third, based on the framework, a trained model can be regarded as representative of a group’s behaviour, which can then be interrogated in control conditions using off-policy simulations to gain insights into the learning processes behind subject’s choices. Finally, the framework can be used to predict the diagnostic labels of the subjects.

It might be possible to design different variants of *Q*-learning models (e.g., based on the analysis presented above) and obtain more competitive prediction accuracy. For example, although it is non-trivial, it is possible to design a new variant of gql able to track oscillatory behaviour such as that described here. Our aim was not to rule out this possibility, but rather to show that the framework can automatically extract learning features from subjects’ actions using learning to learn principles without requiring feature engineering in the models. This is even when those features were initially invisible in task performance metrics.

Our approach inherits these benefits from the field of neural networks, in which feature engineering has been significantly simplified across various domains (Lecun et al., 2015). However, our approach also inherits the black-box nature of these neural networks, i.e., the lack of an interpretable working mechanism. This might not be an issue in some applications, such as the ones mentioned above; however, this needs to be addressed in other applications in which the aim of the study is actually to obtaining an interpretable working mechanism. Nevertheless, we were able to show that running controlled experiments on the model using off-policy simulations can provide significant insights into the processes that mediate subjects’ choices. Interpreting neural networks is an active area of research in machine learning (e.g., Karpathy et al., 2015), and the approach proposed here will benefit from further developments in this area.

In particular, although we found that off-policy simulations of the model could be used to gain insights into the model’s working mechanism, off-policy simulations need to be designed manually to determine inputs to the model. Here, we designed the initial off-policy simulations based on the specific questions and hypotheses that we were interested in testing and using overall behavioural statistics (Figure 6; Section S2). However, an important aspect of the behavioural process, i.e., the tendency of subjects to oscillate between the actions, was not visible in those simulations and, because of this, we had to design another set of inputs to investigate these oscillations (Figure 10). This shows that the choice of off-policy simulation can affect the interpretation of the model’s working mechanism. As such, although rnn can be trained automatically and without intuition into the behavioural processes behind actions (e.g., Barak, 2017), designing off-policy simulations is not automated and requires manual hypothesis generation. Automating this process will require a method that generates representative inputs (and network outputs) that are clearly able to discriminate the differences between the psychiatric groups. The existence of adversarial examples in neural networks (Szegedy et al., 2013) suggests that this will not be as simple as using the networks searching explicitly for input sequences that are most discriminative – representativeness is also critical.

Recurrent neural networks have previously been used to study reward-related decision-making (Song et al., 2017; Zhang et al., 2018), perceptual decision-making, performance in cognitive tasks, working-memory (Miconi, 2017; Carnevale et al., 2015; Mante et al., 2013; Song et al., 2016; Barak et al., 2013; Yang et al., 2017), motor patterns, motor reach and timing (Sussillo et al., 2015; Hennequin et al., 2014; Rajan et al., 2016; Laje and Buonomano, 2013). Typically, in these studies, an rnn is itself trained to perform the task. This is different from the current study, in which the aim of training was to generate behaviour similar to the subjects’, even if that were to lead to poor performance on the task. One exception is the study of Sussillo et al. (2015) in which a network was trained to generate outputs similar to electromyographic (EMG) signals recorded in behaving animals during a motor reach task. Interestingly, that study found that, even though the model was trained based purely on EMG signals, the internal activity of the model resembled the neural responses recorded from the subjects’ motor cortex. A similar approach could be employed in future research to investigate whether brain activity during decision-making is related to network activity.

With regard to predicting subjects’ diagnostic labels, it was perhaps not surprising to find that the model was unable to achieve a high level of classification accuracy. This is because there is a high level of heterogeneity in patients with the same diagnostic label. Heterogeneity, which is well understood in the wide variation in treatments and treatment outcomes in disorders like depression (e.g., Rush et al., 2006), is likely also to be reflected in differing learning and choice abilities of the subjects.

In the model fitting procedure used here, a single model was fitted to all of the subjects in each group, despite possible individual differences within a group. This was partly because we were interested in obtaining a single parameter set for making predictions for the subject withheld in the leave-one-out cross-validation experiments. Even if a mixed-effect model was fitted to the data, a summary of group statistics will be required for making predictions about a new subject. In other applications, one might be interested in estimating parameters for each individual (either network weights or parameters of the reinforcement-learning models); in this respect using a hierarchical, model fitting procedure would be a more appropriate approach, something that has been done previously for reinforcement-learning models (e.g., Piray et al., 2014) and would be an interesting future step for rnn models.

Along the same lines, due to its rich set of parameters, a single rnn model might be able to learn about and detect individual differences (e.g., differences in the learning-rates of subjects) at an early stage of the task, and then use this information for making predictions about performance on later trials. For example, in the learning-to-learn phase, the model might learn that subjects have either a very high, or a very low learning-rate. Then, when being evaluated in the actual learning task, the model can use observations from subjects’ choices on early trials to determine whether the learning-rate for that specific subject is high or low, and then utilise that information for making more accurate predictions in latter trials. Determining individual-specific traits in early trials of the task is presumably *not part of the computational processes* occurring in the subject’s brain during the task, but it is occurring in the model merely to make more accurate predictions. To the extent that the network learns such higher order structure, it is appealing, though hard, to extract information about the heterogeneity from the recurrent state of the rnn. Of course, it implies that the (implicit) inferences that the rnn makes about the type of subject might be confounded with the (implicit) inferences that the rnn makes about the actual choices – thus it is a model that makes predictions about subjects’ choices using mechanisms that may not be competent computational models of the way that the subjects themselves make those choices.

## Materials and methods

### Participants

34 uni-polar depression (depression), 33 bipolar (bipolar) and 34 control (healthy) participants (age, gender, IQ and education matched) were recruited from outpatient mental health clinics at the Brain and Mind Research Institute, Sydney, and the surrounding community. Participants were aged between 16 and 33 years. Exclusion criteria for both clinical and control groups were history of neurological disease (e.g. head trauma, epilepsy), medical illness known to impact cognitive and brain function (e.g. cancer), intellectual and/or developmental disability and insufficient English for neuropsychological assessment. Controls were screened for psychopathology by a research psychologist via clinical interview. Patients were tested under ’treatment-as-usual’ conditions, and at the time of assessment, 77% of depressed and 85% of bipolar patients were taking medications (see Table 2 for breakdown of medication use). The study was approved by the University of Sydney ethics committee. Participants gave informed consent prior to participation in the study.

Demographics and clinical characteristics of the sample are presented in Table 2. Levene’s test indicated unequal variances for the HDRS (Hamilton Depression Rating Scale; Hamilton, 1960), YMRS (Young Mania Rating Scale; Young et al., 1978), SOFAS (Social and Occupational Functional Scale; Goldman et al., 1992) and age, thus Welch’s statistic was used for these variables. A one-way ANOVA revealed no differences between groups in age [*F* (2, 98) = 2.48, *p* = 0.09], education [*F* (2, 98) = 1.76, *p* = 0.18], IQ [*F* (2, 94) = 0.47, *p* = 0.62] or gender (χ^2^ = 2.66, *p* = 0.27). There were differences in HDRS [*F* (2, 49.21) = 64.21, *p*< 0.001], YMRS [*F* (2, 43.71) = 12.57, *p*< 0.001], and SOFAS [*F* (2, 41.61) = 169.66, *p*< 0.001]. Bonferroni post-hoc comparisons revealed higher depression scores in depression group compared to bipolar and healthy groups, and higher depression in bipolar group compared to healthy group. Mania scores were significantly higher in the bipolar group compared to the healthy group. Both patient groups had significantly lower SOFAS scores compared to the healthy group, but did not differ from one another. Age of mental illness onset was younger in the depression group compared to the bipolar group [*t*(56) = −2.14, *p* = 0.04], however duration of illness did not differ significantly between groups [*t*(56) = 1.25, *p* = 0.22]. There were no differences between groups in pre-test hunger [*F* (2, 79) = 0.54, *p* = 0.59] or average snack rating [*F* (2, 79) = 2.53, *p* = 0.09].

### Task

The instrumental learning task (Figure 2) involved participants choosing between pressing a left or right button in order to earn food rewards (an M&M chocolate or a BBQ flavoured cracker). We refer to these two key presses as L and R for left and right button presses respectively. Fourteen healthy participants (41.2% of the group) and 13 bipolar participants (36.7% of the group) completed the task in an fMRI setting, using a 2 button Lumina response box. The remaining healthy and bipolar participants, and all depression participants, completed the task on a computer with a keyboard, where the “Z” and “?” keys were designated L and R. Although the performance of subjects was higher overall in the fMRI setting [*β* = 0.050, SE=0.024, *p* = 0.041], the place in which the task was completed had no significant effect on how choices adjusted on a trial-by-trial basis, either on the probability of staying with the same action after earning a reward [*β* = 0.041, SE=0.054, *p* = 0.45]^12^, or after no reward [*β* = 0.030, *SE*=0.062, *p* = 0.627]), and, therefore, the data were combined.

During each block, one action was always associated with a higher probability of reward than the other. Across blocks, the action with the higher reward probability switched identities (left or right), and the probabilities varied between 0.25, 0.125, and 0.08. The probability of reward on the other action always remained at 0.05. Participants were instructed to earn as many points as possible, as they would be given the concomitant number of M&Ms or BBQ flavoured crackers at the end of the session. After a non-rewarded response, a grey circle appeared in the centre of the screen for 250ms, whereas after a rewarded response the key turned green and an image of the food reward earned appeared in the centre of the screen for 500ms. A tally of accumulated winnings remained on the bottom of the screen for the duration of the task. The task began with a 0.25 contingency practice block and a pleasantness rating for each food outcome (−5 to +5). Responding was self-paced during the 12 blocks of training, each 40-s in length. During inter-block intervals (12 seconds) the participants rated how causal each button was in earning rewards. These self-reports (causal ratings) are not used in the modelling analysis presented here.

**Table 2.** Demographic and clinical characteristics of participants. Means(SD). HDRS: Hamilton Depression Rating Scale; YMRS: Young Mania Rating Scale; SOFAS: Social and Occupational Functioning Scale; a: depression greater than healthy and bipolar, *p*< 0.05. b: bipolar greater than healthy, *p*< 0.05. c: healthy greater than depression and bipolar, *p*< 0.05. d: depression slower than healthy and bipolar, *p*< 0.05.

|  | HEALTHY<br>(n=34) | DEPRESSION<br>(n=34) | BIPOLAR<br>(n=33) |
| --- | --- | --- | --- |
| Demographics |  |  |  |
| Gender (M:F) | 15:19 | 15:19 | 9:24 |
| Age in years | 23.6 (4.3) | 21.6 (2.5) | 23.1 (4.4) |
| Predicted IQ | 107.3 (7.5) | 105.5 (7.9) | 106.0 (7.4) |
| Education | 14.3 (3.0) | 13.3(1.9) | 13.3 (2.4) |
| Symptoms and History |  |  |  |
| Age of onset (years) | - | 14.4 (3.8) | 15.9 (4.7) |
| Duration of illness (years) | - | 7.7 (4.3) | 6.4 (3.3) |
| HDRS | 1.5(2.0) | 14.1 (7.2)<sup>a</sup> | 8.9 (6.5) |
| YMRS | 0.1 (0.4) | 2.5 (5.4) | 4.6 (5.8)<sup>b</sup> |
| SOFAS | 91.0 (3.5)<sup>c</sup> | 63.8 (9.2) | 65.7 (13.7) |
| Medication |  |  |  |
| Medicated | - | 77% | 85% |
| Anti-depressants | - | 71% | 41% |
| Mood stabilizers/Anti-convulsants | - | 9% | 73% |
| Lithium | - | 0% | 18% |
| Anti-psychotics | - | 18% | 33% |
| Anxiolytics | - | 0% | 3% |
| Motivation measures |  |  |  |
| Hunger | 6.5 (1.7) | 6.0 (2.1) | 6.0 (2.4) |
| Reward Pleasantness | 3.1 (1.3) | 2.0 (2.0) | 2.6 (2.0) |
| Press Rate/Sec | 0.93 (0.3) | 0.71 (0.2)<sup>d</sup> | 0.98 (0.3) |

Duration of illness indicates time since patient first experienced mental health problems, not time since diagnosis.

### Computational models

#### Notation

The set of available actions is denoted by 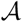. Here 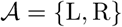, with L and R referring to left and right key presses respectively. A set of subjects is denoted by 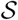, and the total number of trials completed by subject 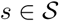 over the whole task (all blocks) is denoted by 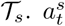 adenotes the action taken by subject *s* at trial *t*. The reward earned at trial *t* is denoted by *r_t_*, and we use *a_t_* to refer to an action taken at time *t*, either by the subjects or the models (in simulations).

#### Recurrent neural network model (rnn)

The architecture used is based on recurrent neural network model (rnn) and is depicted in Figure 1. The model is composed of an lstm layer (Long short-term memory; Hochreiter and Schmidhuber, 1997) and an output softmax layer with two nodes (since there are two actions in the task). The inputs to the lstm layer are the previous action (*a_t_*_−1_ coded using one-hot transformation) and the reward received after taking action (*r_t_*_−1_ ∈ {0, 1}). The outputs of the softmax are probabilities of selecting each action, which are denoted by *π_t_*(*a*; rnn) for action 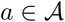 at trial *t*.

In the learning-to-learn phase, the aim is to train weights in the network so that the model learns to predict subjects’ actions given their past observations (i.e., it learns how *they* learn). For this purpose, the objective function for optimising weights in the network (denoted by Θ) for subject set 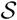 is,

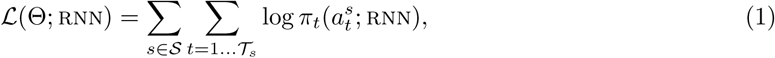
where 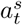 is the action selected by subjects *s* at trial *t*, and *π_t_*(.; rnn) is the probability that model assigns to each action. Note that the policy is conditioned on the previous actions and rewards in each block of training; notation for this is omitted, for simplicity.

Models were trained using the maximum-likelihood (ML) estimation method,

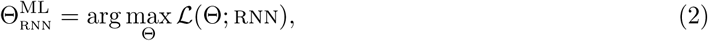
where Θ is a vector containing free-parameters of the model (in both lstm and softmax layers). The models were implemented in TensorFlow (Abadi et al., 2016) and optimized using Adam optimizer (Kingma and Ba, 2014). Note that Θ was estimated for each group of subjects separately. Networks with different numbers *N_c_* of lstm cells (*N_c_* ∈ {5, 10, 20}) were considered, and the best model was selected using leave-one-out cross-validation (see below). Early stopping was used for regularization and the optimal number of training iterations was selected using leave-one-out cross-validation.

The total number of free parameters (in both the lstm layer and softmax layer) were 190, 580, and 1960 for the networks with 5, 10, and 20 lstm cells, respectively. In order to control for the effect of initialization of network weights on the final results, a single random network of each size (5, 10, 20) was generated, and was used to initialize the weights in the network.

After the learning-to-learn phase, the weights in the network were frozen and the trained model was used for three purposes: (i) cross-validation (see below), (ii) on-policy simulations and (iii) off-policy simulations. For cross-validation, the previous actions of the test subject(s) and the rewards experienced by the subject(s) were fed into the model, but unlike the learning-to-learn phase, the weights were not changing and we only recorded the prediction of the model about the next action. Note that even though the weights in the network were fixed, the output of the network changed from trial to trial due to the recurrent nature of these networks. Also, due to the small sample size, we used the same set of subjects for testing the model and for the validation of model hyper-parameters (*N_c_* and number of optimization iterations).

Other than being used for calculating cross-validation statistics, trained models were used for on-policy and off-policy simulations (with frozen weights). In the on-policy simulations, the model received its own actions and earned rewards as inputs (instead of receiving the action selected by the subjects). In the off-policy simulations, the set of actions and rewards that the model received was fixed and predetermined. The details of these simulations are reported in the Results section.

**Model settings**. For the rnn model, leave-one-out cross-validation was used to determine the number of cells and optimisation iterations required for the rnn model to achieve the highest prediction accuracy. We found that the lowest mean negative log-probability (nlp) was achieved by 10 cells in the LSMT layer and after 1100, 1200 optimisation iterations for the healthy and depression groups respectively whereas for the bipolar group the best nlp was achieved by 20 cells and 400 optimisation iterations (see Figure S1). These settings were used for making predictions and simulations.

##### Baseline methods

We used three baseline methods, ql, qlp and gql, which are variants and generalizations of *Q*-learning (Watkins, 1989).

**ql model**. After taking action *a_t_*_−1_ at time *t* − 1, the value of the action, denoted by *Q_t_*(*a_t_*_−1_), is updated as follows,

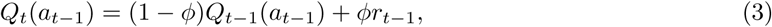
where *ϕ* is the learning-rate and *r_t_*_−1_ is the reward received after taking the action. Given the action values, the probability of taking action *a* ∈ {L, R} in trial *t* is:

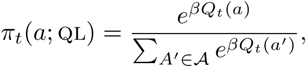
where *β* > 0 is a free-parameter and controls the contribution of values to the choices (balance between exploration and exploitation). The free-parameters of this variant are *ϕ* and *β*. Note that the probability that the models predict for each action at trial *t* is necessarily based on the data *before* observing the action and reward at trial *t*. Further, since there are only two actions, we can write *π_t_*(L; ql)=1 − *π_t_*(R; ql)= *σ*(*β*(*Q_t_*(L) − *Q_t_*(R))) where *σ*(·) is the standard logistic sigmoid.

**qlp model**. This model is inspired by the fact that humans and other animals have a tendency to stick with the same action for multiple trials (i.e., perseverate), or sometimes to alternate between the actions (independent of the reward effects; Lau and Glimcher, 2005). We therefore call this model qlp, for *Q*-learning with perseveration. In it, action values are updated according to equation 3 and so similarly to the ql model, but the probability of selecting actions is,

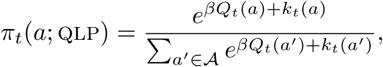
where,

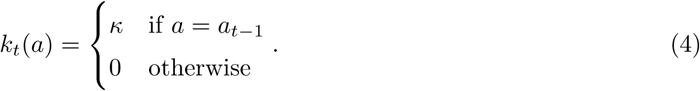

Therefore, there is a tendency to select the same action again on the next trial (if *κ* > 0) or switch to the other action (if *κ* < 0). In the specific case that *κ* = 0, the qlp model reduces to ql. Free-parameters are *ϕ, β, κ*.

**gql model**. As we will show in the results section, neither ql nor qlp fit the behaviour of the subjects in the task. As such, we aimed to develop a baseline model which could at least capture high-level behavioural trends, and we built a generalised *Q*-learning model, gql, to compare with rnn. In this variant, instead of learning a single action value for each action, the model learns *N* different values for each action, where the difference between the values learned for each action is that they are updated using different learning-rates. The action values for action *a* are denoted by ***Q***(*a*), which is a vector of size *N*, and the corresponding learning-rates are denoted by vector Φ of size 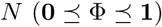. Based on this, the value of action *a_t_*_−1_ at trial *t* − 1 is updated as follows,

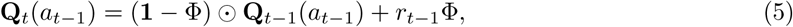
where ☉ represents the element-wise Hadamard product. For example, if *N* = 2, and Φ = [0.1, 0.05], then the model will learn two different values for each action (L, R actions) with one of the values updated using a learning-rate of 0.1 and the other updated using a learning-rate of 0.05. In the specific case that *N* = 1, the above equation reduces to equation 3 used in ql and qlp models, in which only a single value is learned for each action.

In the qlp model, the current action is affected by the last taken action (perseveration). This property is generalised in the gql model by learning the history of previously taken actions instead of just the last action. These action histories are denoted by **H**(*a*) for action *a*. **H**(*a*) is a vector of size *N*, and each entry of this vector tracks the tendency of taking action *a* in the past, i.e., if an element of **H**(*a*) is close to one it means that action *a* was taken frequently in the past and being close to zero implies that the action was taken rarely. In similar fashion to action values, for each action *N* different histories are tracked, each of which is modulated by a separate learning-rate. Learning-rates are represented in vector Ψ of size 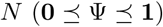. Assuming that action *a_t_*_−1_ was taken at trial *t* − 1, **H**(*a*) updates as follows,

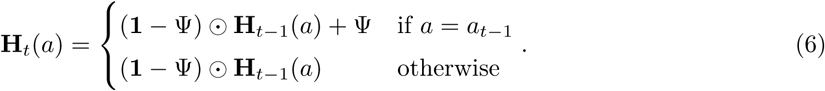

Intuitively, according to the above equation, if action a was taken on a trial, **H**(*a*) increases (the amount of increase depends on the learning-rate of each entry), and for the rest of the actions, **H**(other actions) will decrease (again the amount of decrement is modulated by the learning rates). For example, if *N* = 2, and Ψ = [0.1, 0.05], it means that for each action two choice tendencies will be learned, one of which is updated by rate 0.1 and the other one by rate 0.05.

Having learned *Q*(*a*) and **H**(*a*) for each action, the next question is how are they combined to guide choice. *Q*-learning models assume that the contribution of values to choices is modulated by parameter *β*. Here, since the model learns multiple values for each action, we assume that each value is weighted by a separate parameter, denoted by vector **B** of size *N*. Similarly, in the qlp model the contribution of perseveration to choices is controlled by parameter *κ*, and here we assume that parameter **K** modulates the contribution of previous actions to the current choice. Based on this, the probability of taking action *a* at trial *t* is,

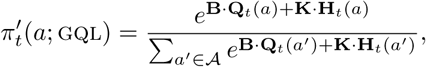
where “·” operator refers to the inner product. Here, we also add extra flexibility to the model by allowing values to interact with the history of previous actions in influencing choices. For example, if *N* = 2, we allow the two learned values for each action to interact with the two learned action histories of each action, leading to four interaction terms, and the contribution of each interaction term to choices is determined by a matrix C of size *N* × *N* (*N* = 2 in this example),

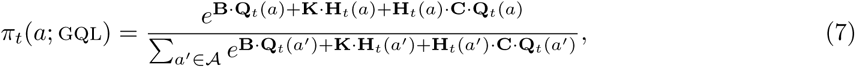

The free-parameters of this model are Φ, Ψ, **B, K**, and **C**. In this paper we use models with *N* = 1, 2, 10, which have 5, 12 and 140 free parameters respectively. We used *N* = 2 for the results reported in the main text, since this model setting was able to capture several behavioural trends while still being interpretable. The results using *N* = 1, 10 are reported in the supplementary materials to illustrate the models’ capabilities in extreme cases.

**Objective function**. The objective function for optimising the models was the same as the one chosen for rnn,

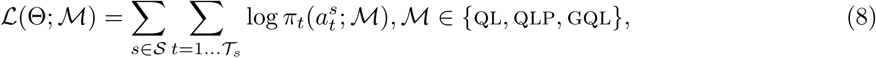
where, as mentioned before, 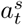 is the action selected by subject *s* at trial *t*, and 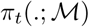 is the probability that model 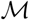 assigns to each action. Models were trained using the maximum-likelihood estimation method,

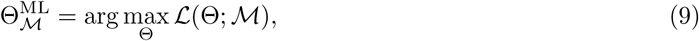
where Θ is a vector containing the free-parameters of the models. Optimizations for all models were performed using Adam optimizer (Kingma and Ba, 2014), and using the automatic differentiation method provided in TensorFlow (Abadi et al., 2016). The free-parameters with limited support (*ϕ, β*, Φ, Ψ) were transformed to satisfy the constraints.

### Performance measures

Two different measures were used for quantifying the predictive accuracy of the models. The first measure is the average log-probability of the models’ prediction for the actions taken by subjects. For a group of subjects denoted by 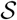, we define negative log-probability (nlp) as follows:

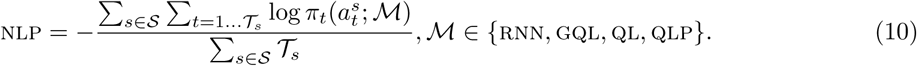

The other measure is the percentage of actions predicted correctly,

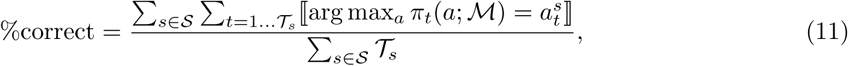
where 〚.〛denotes the indicator function. Unlike, ‘%correct’, nlp takes the probabilities of predictions into account instead of making binary predictions for the next action. In this way, if the models are certain about wrong predictions nlp performance gets penalized, and it gets credit if the models are certain about a correct prediction.

### Model selection

Leave-one-out cross-validation was used for comparing different models. At each round, one of the subjects was withheld and the model was trained using the remaining subjects; the trained model was then used to make predictions about the withheld subject. The withheld subject was rotated in each group, yielding 34, 34 and 33 prediction accuracy measures in the healthy, depression, and bipolar groups respectively.

### Statistical analysis

For the analysis we performed hierarchical linear mixed-effects regression using the lme4 package in R (Bates et al., 2015) and obtained *p*-values for regression coefficients using the lmerTest package (Kuznetsova et al., 2016). For each test we report parameter estimate (*β*), standard error (SE), and *p*-value.

## Acknowledgments

This research was supported by a grant from the NHMRC, GNT1089270. BB was supported by a Senior Principal Research Fellowship from the NHMRC, GNT1079561. PD was partly funded by the Gatsby Charitable Foundation. PD is currently on a leave of absence at Uber Technologies. None of these bodies influenced this study.

## Supporting information

### S1 Behavioural analysis using gql

In the gql simulations presented in Figure 6, we observed that earning a reward (shown by black crosses) causes a ‘dip’ in the probability of staying with an action, but that the dip decreases in magnitude with the number of recent rewards. Neither of these phenomena arises from ql and qlp; gql captures them because it learns that the values of each action are updated at two different rates, positively with reward at a slow rate (0.145); and negatively with reward a fast rate (0.815; see Table S4).

The dip implies a tendency to switch to the other action. This is consistent with a phenomenon that is apparent in Figure 4, namely that the probability of switching increases after rewards. The dip is produced by the fast updating process with its negative reward contribution.

However, the fast process is also fast to forget, and so has an influence that cannot accumulate. The slow process ultimately exerts a stronger effect; and furthermore accumulates over multiple rewards in a way that the fast process cannot accomplish.

Based on this, allowing the model to track two different values for each action is important, and the model will not be able to produce this behaviour if it tracks only one value for each action (*N* = 1) as shown in Figure S8.

### S2 The choice of off-policy settings

In the simulations shown in Figure 6, action R is fed into the model for the first 10 trials; then a switch is made to action L. This is based on the fact that in the empirical data, the average length of staying with an action (when one reward is earned in the middle of the ’run’ of the action) is 9.8. The first, second and third rewards in Figure 6 are delivered after an action was taken 4, 12, and 17 times respectively. This is based on the fact that in the empirical data, the average number of key-presses in order to earn the first, second and third rewards is 4.07, 11.6, and 17.4 respectively.

In the simulations shown in Figure 10, the reason for adding leading R before oscillations is to show that the models do not oscillate all the time, but only after they are fed with oscillations. Indeed, qlp is in principle able to produce 1-step oscillations (singe-action runs) by assigning a negative weight to the perseveration parameter, i.e., instead of the model having a tendency to stay on the previously selected action, it will have a tendency to switch from this action. However, under this condition the model will keep oscillating between the actions from trial 1, implying that it can only produce runs of length 1 no matter what the length of the previous run of actions was, which is inconsistent with the empirical data presented in Figure 9.

**Figure S1.**
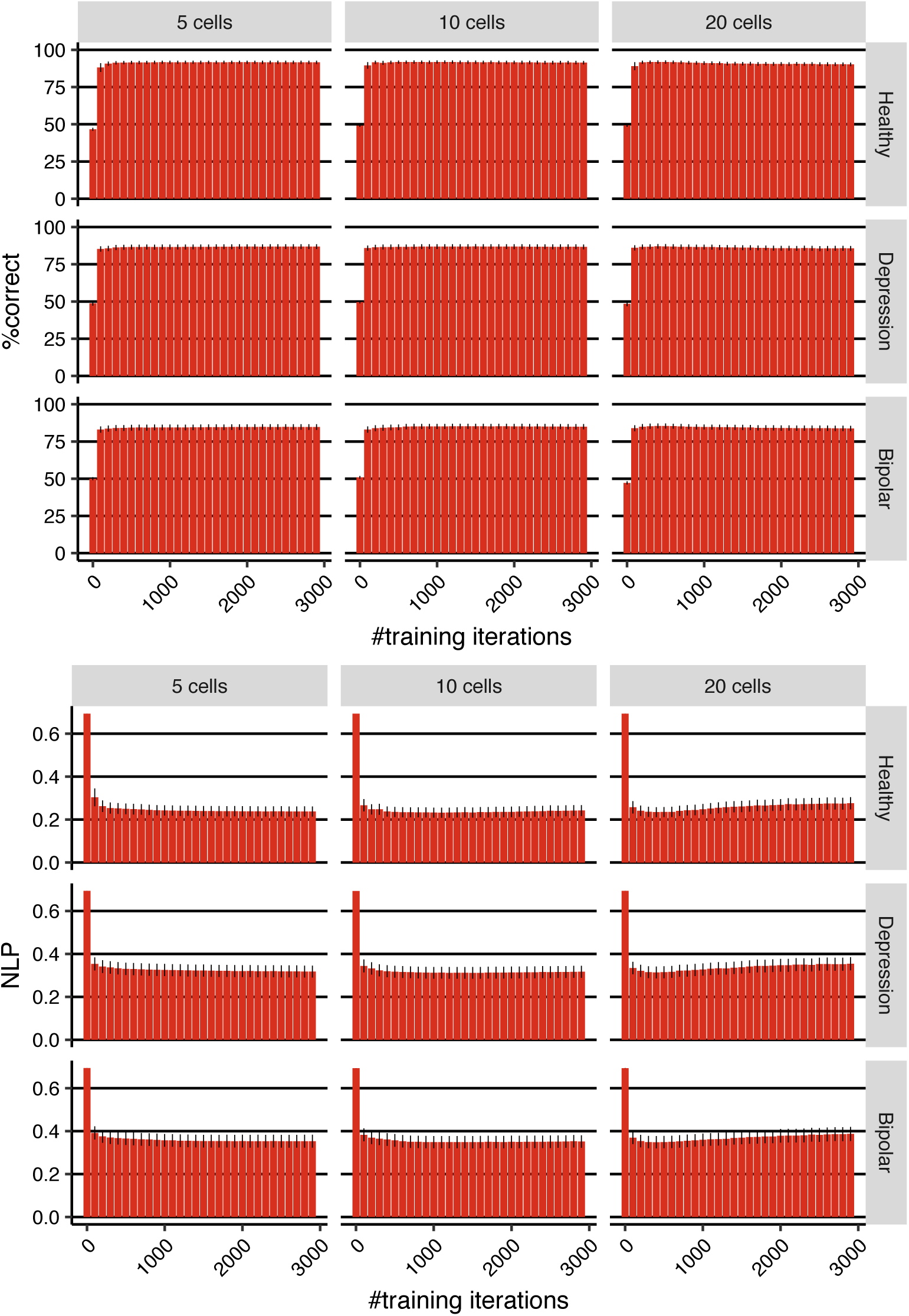
Cross-validation results for different number of cells and optimization iterations. **(Top-panel)** Percentage of actions predicted correctly averaged over leave-one-out cross-validation folds. **(Bottom-panel)** Mean nlp averaged over cross-validation folds. Error-bars represent 1SEM.

**Figure S2.**
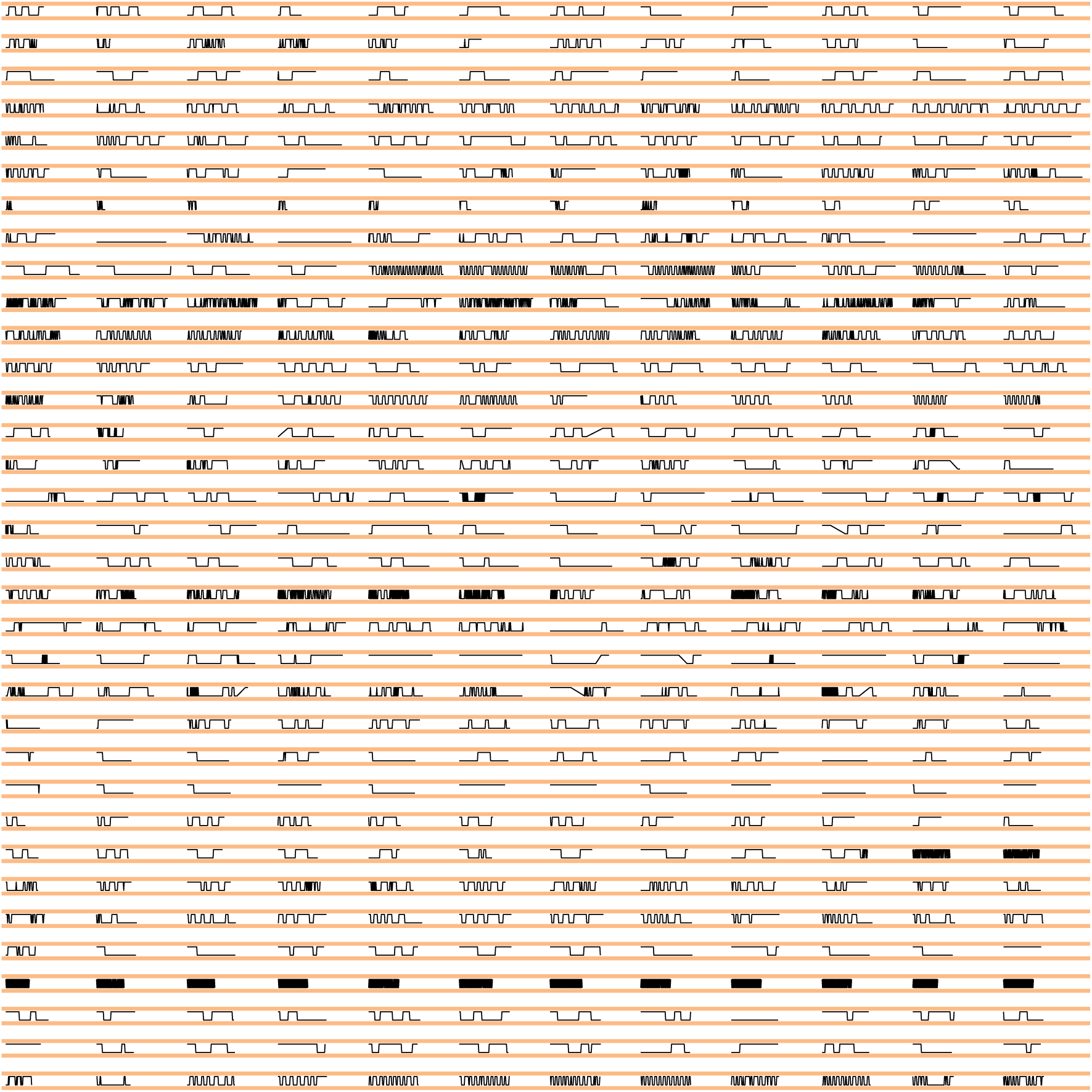
Choices in healthy group. Each row shows choices of a subject across different blocks (12 blocks).

**Figure S3.**
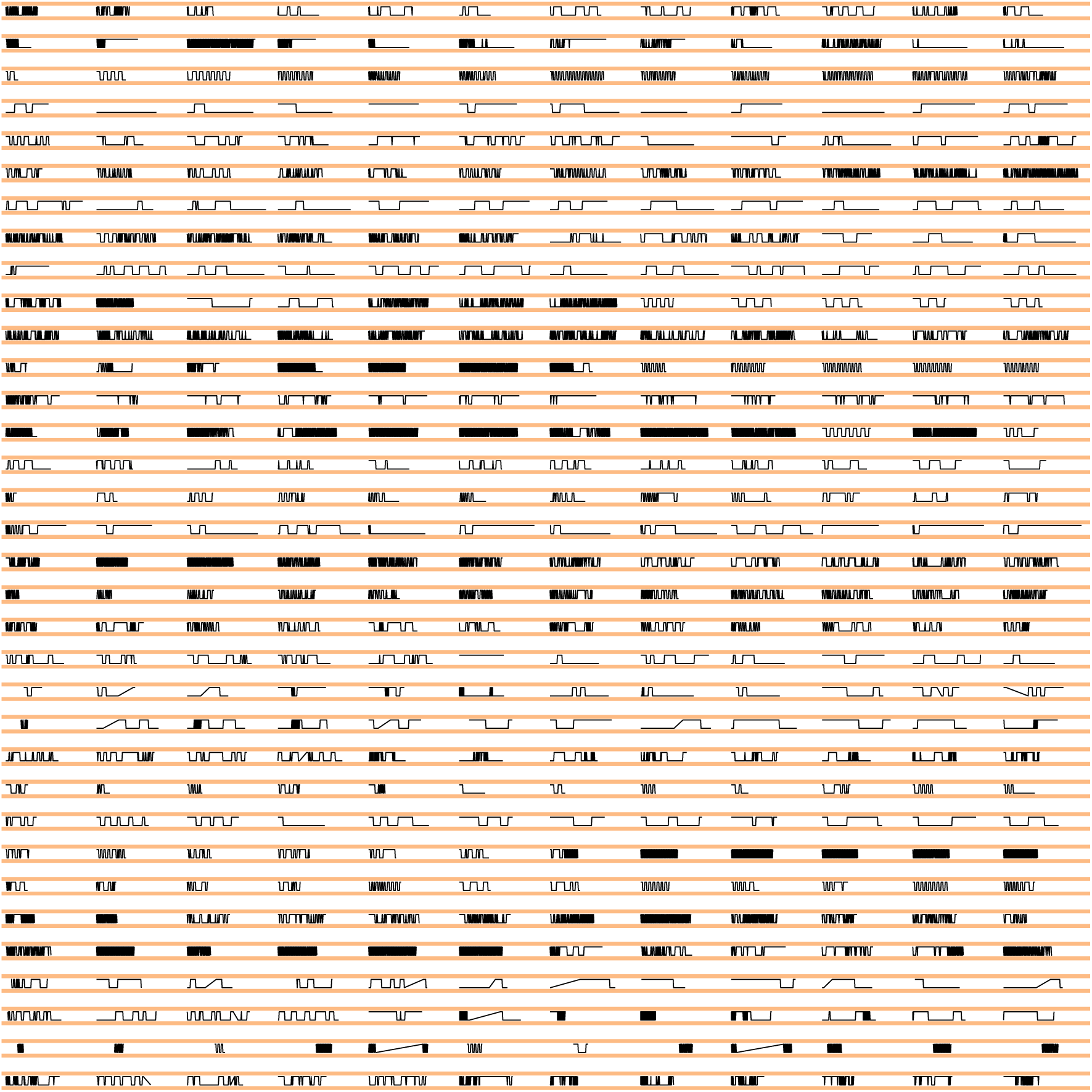
Choices in depression group. Each row shows choices of a subject across different blocks (12 blocks).

**Figure S4.**
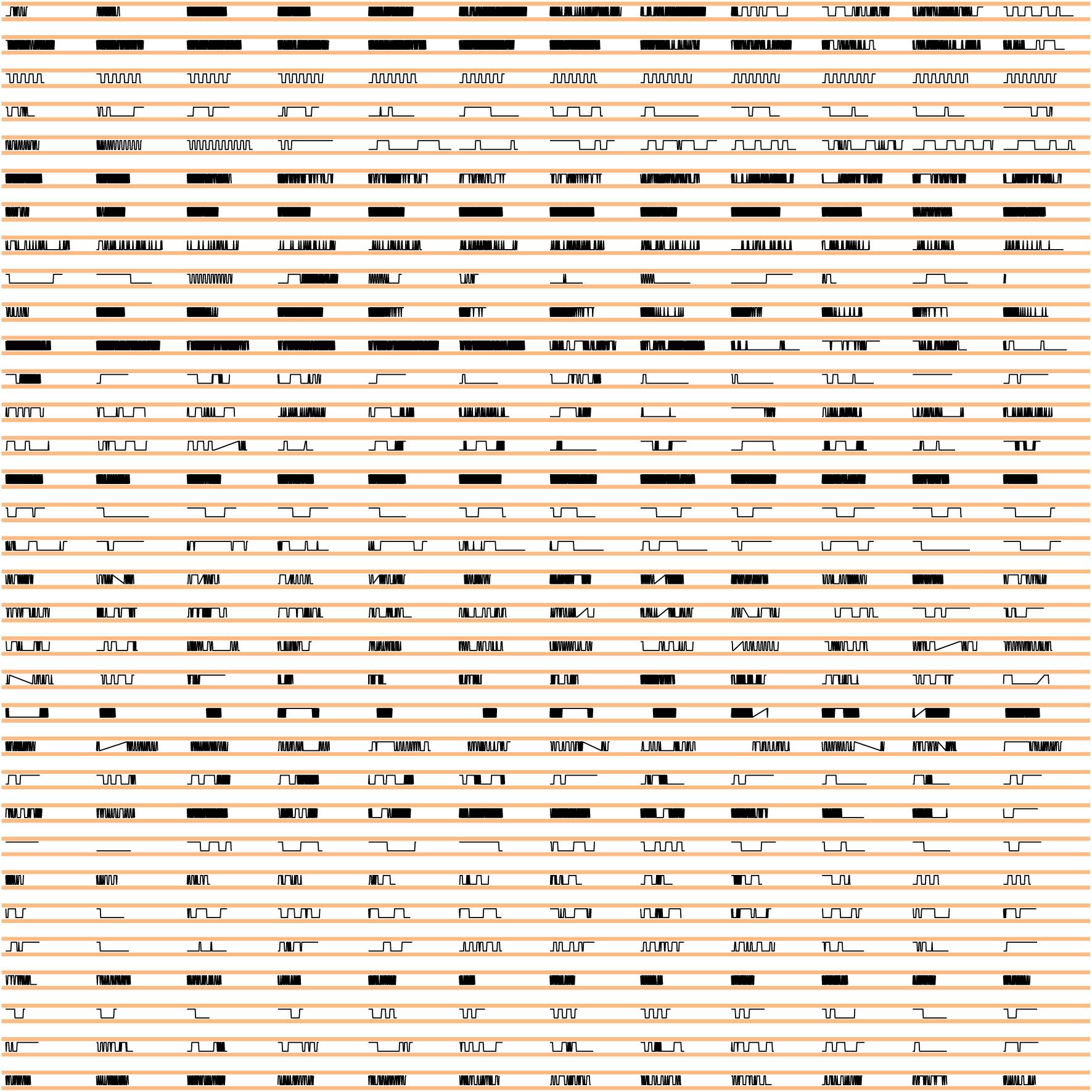
Choices in bipolar group. Each row shows choices of a subject across different blocks (12 blocks).

**Figure S5.**
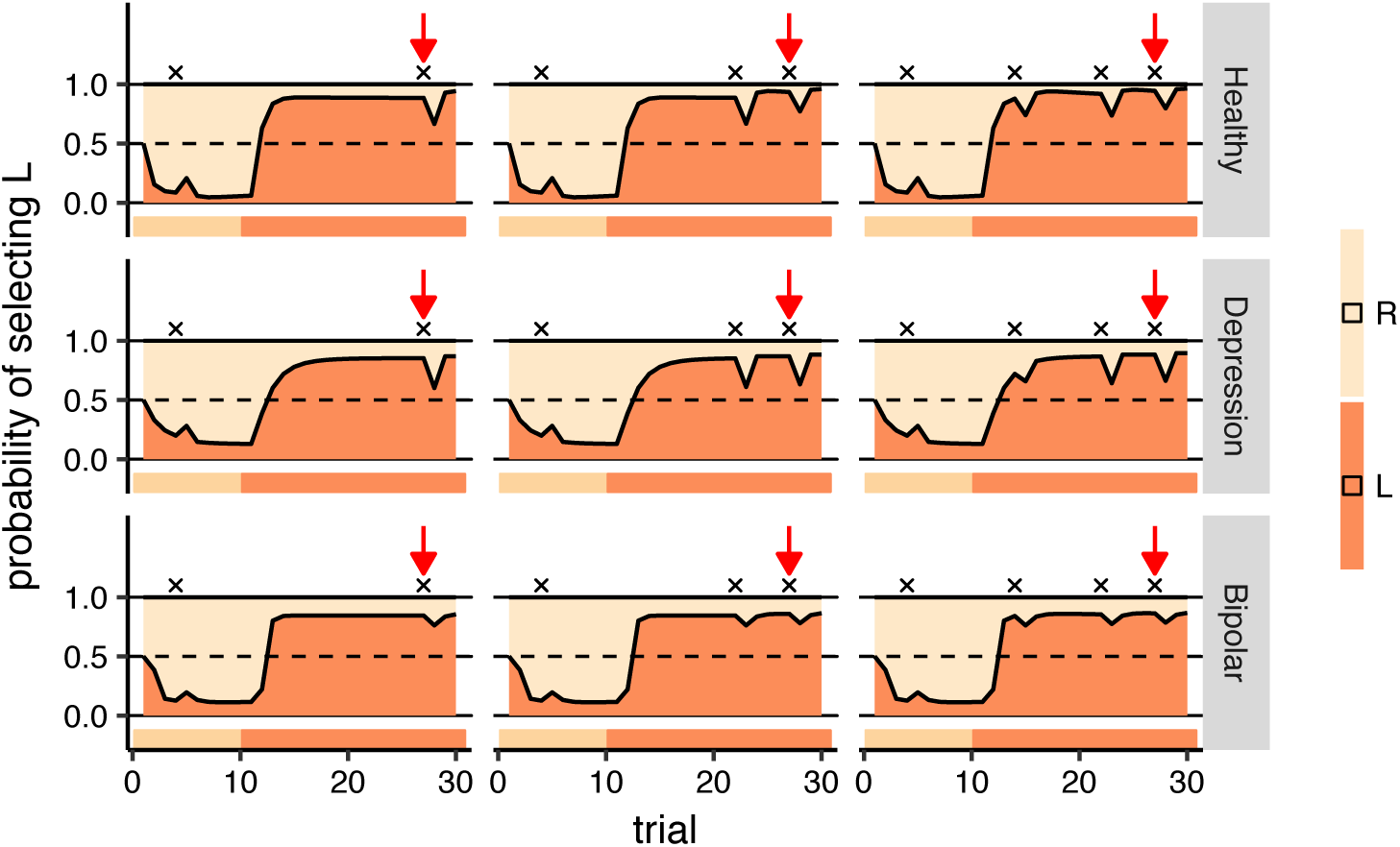
Off-policy simulations of gql (*N* = 2). Each panel shows a simulation for 30 trials (horizontal axis), and the vertical axis shows the predictions of each model on each trial. The ribbon below each panel shows the action which was fed to the model on each trial. In the first 10 trials, the action that the model received was R and in the next 20 trials it was L. Rewards are shown by black crosses (x) on the graphs. See text for the interpretation of the graph. Note that the simulation conditions is same as the one depicted in Figures 7 and 6

**Figure S6.**
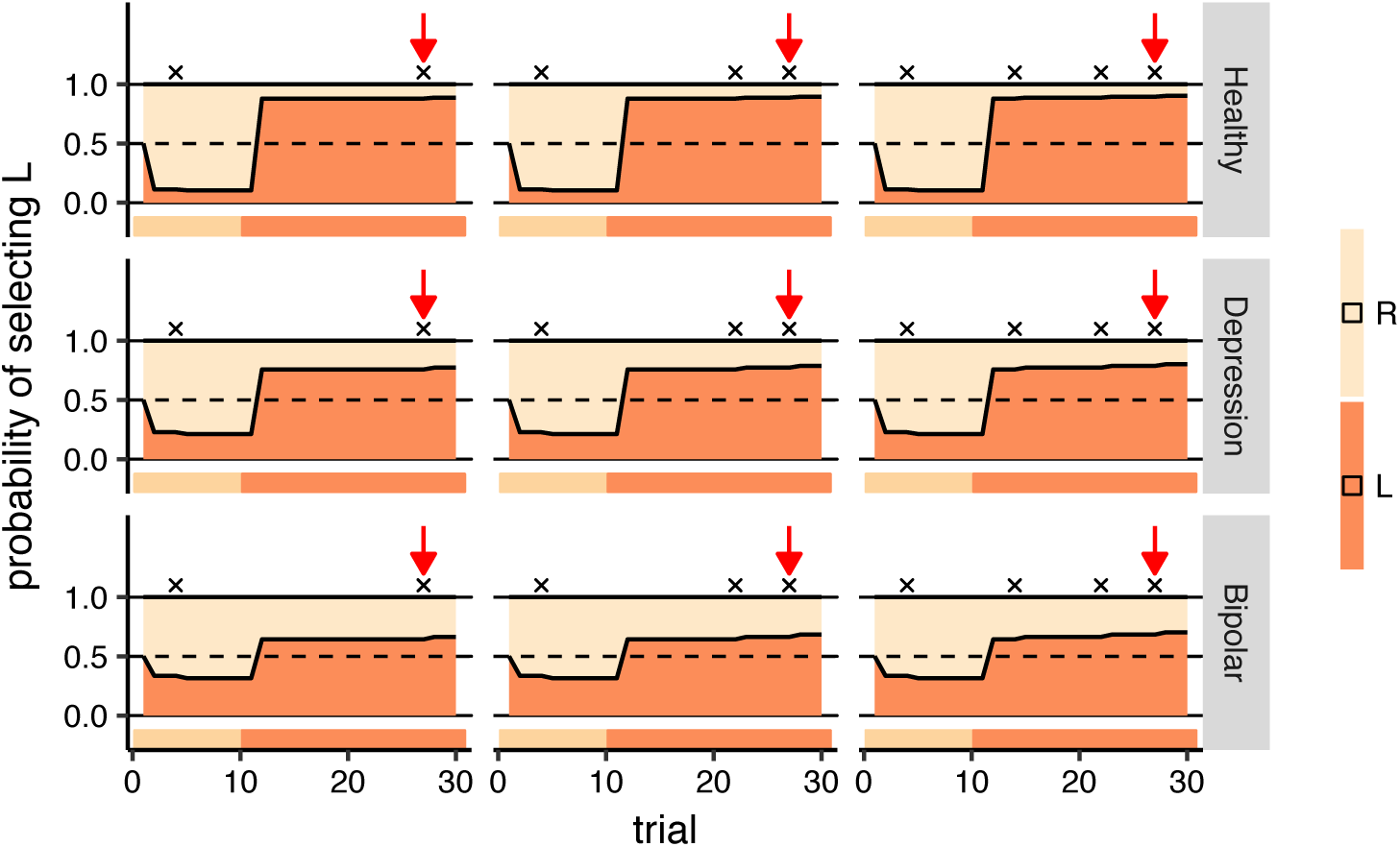
Off-policy simulations of qlp. Each panel shows a simulation for 30 trials (horizontal axis), and the vertical axis shows the predictions of each model on each trial. The ribbon below each panel shows the action which was fed to the model on each trial. In the first 10 trials, the action that the model received was R and in the next 20 trials it was L. Rewards are shown by black crosses (x) on the graphs. See text for the interpretation of the graph. Note that the simulation conditions is same as the one depicted in Figures 7 and 6

**Figure S7.**
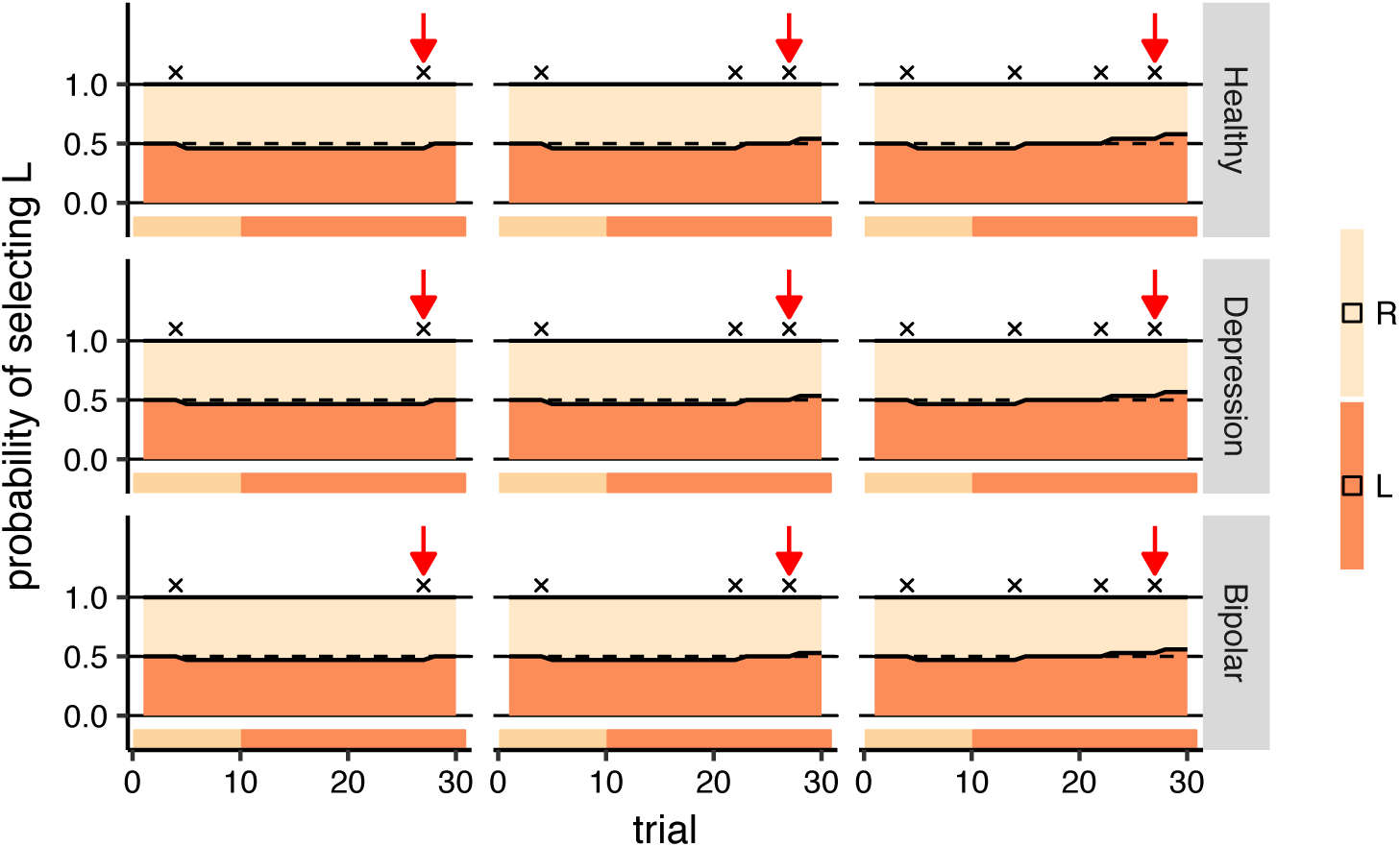
Off-policy simulations of ql. Each panel shows a simulation for 30 trials (horizontal axis), and the vertical axis shows the predictions of each model on each trial. The ribbon below each panel shows the action which was fed to the model on each trial. In the first 10 trials, the action that the model received was R and in the next 20 trials it was L. Rewards are shown by black crosses (x) on the graphs. See text for the interpretation of the graph. Note that the simulation conditions is same as the one depicted in Figures 7 and 6

**Figure S8.**
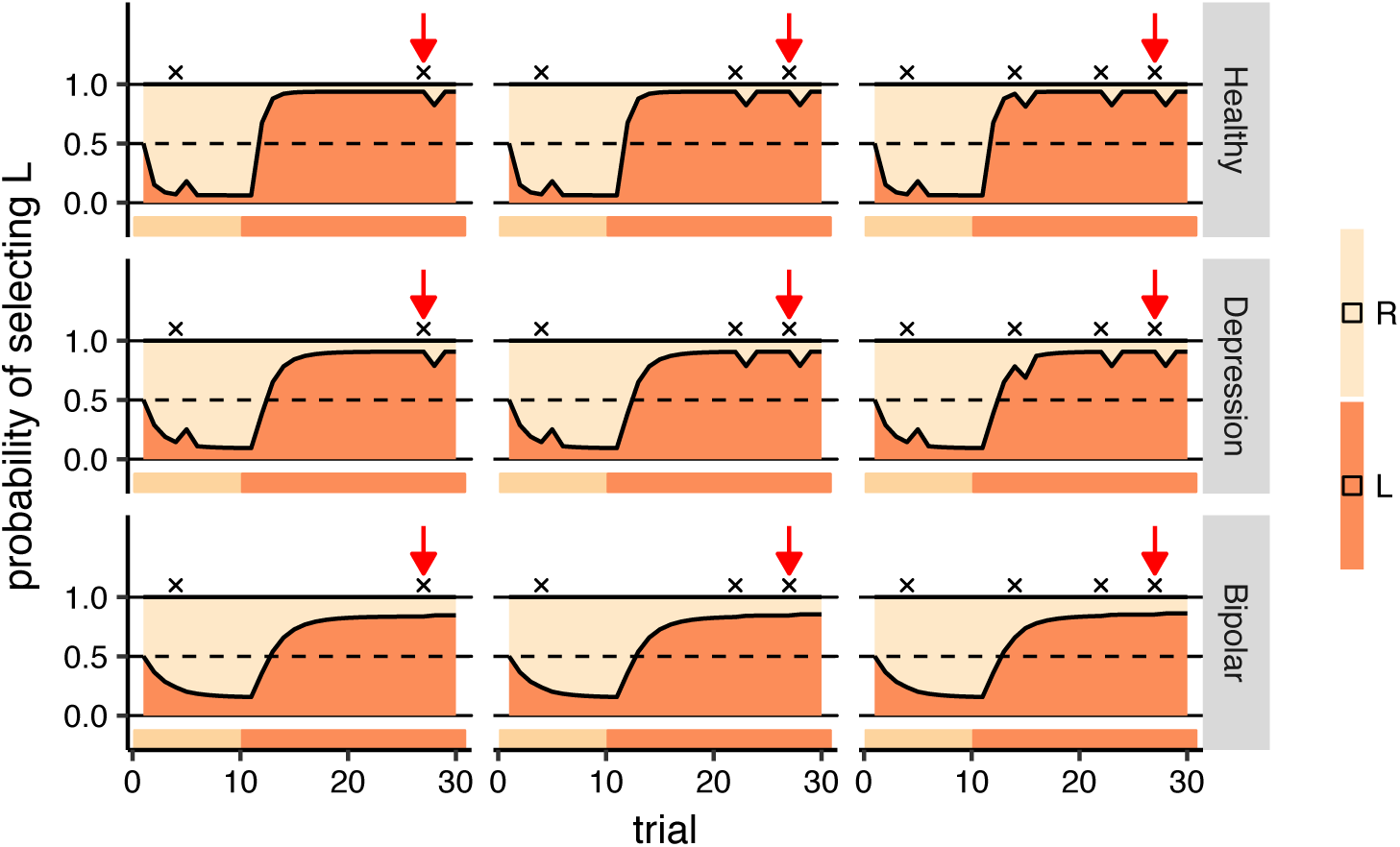
Off-policy simulations of gql with *N* = 1. Each panel shows a simulation for 30 trials (horizontal axis), and the vertical axis shows the predictions of each model on each trial. The ribbon below each panel shows the action which was fed to the model on each trial. In the first 10 trials, the action that the model received was R and in the next 20 trials it was L. Rewards are shown by black crosses (x) on the graphs. See text for the interpretation of the graph. Note that the simulation conditions is same as the one depicted in Figures 7 and 6

**Figure S9.**
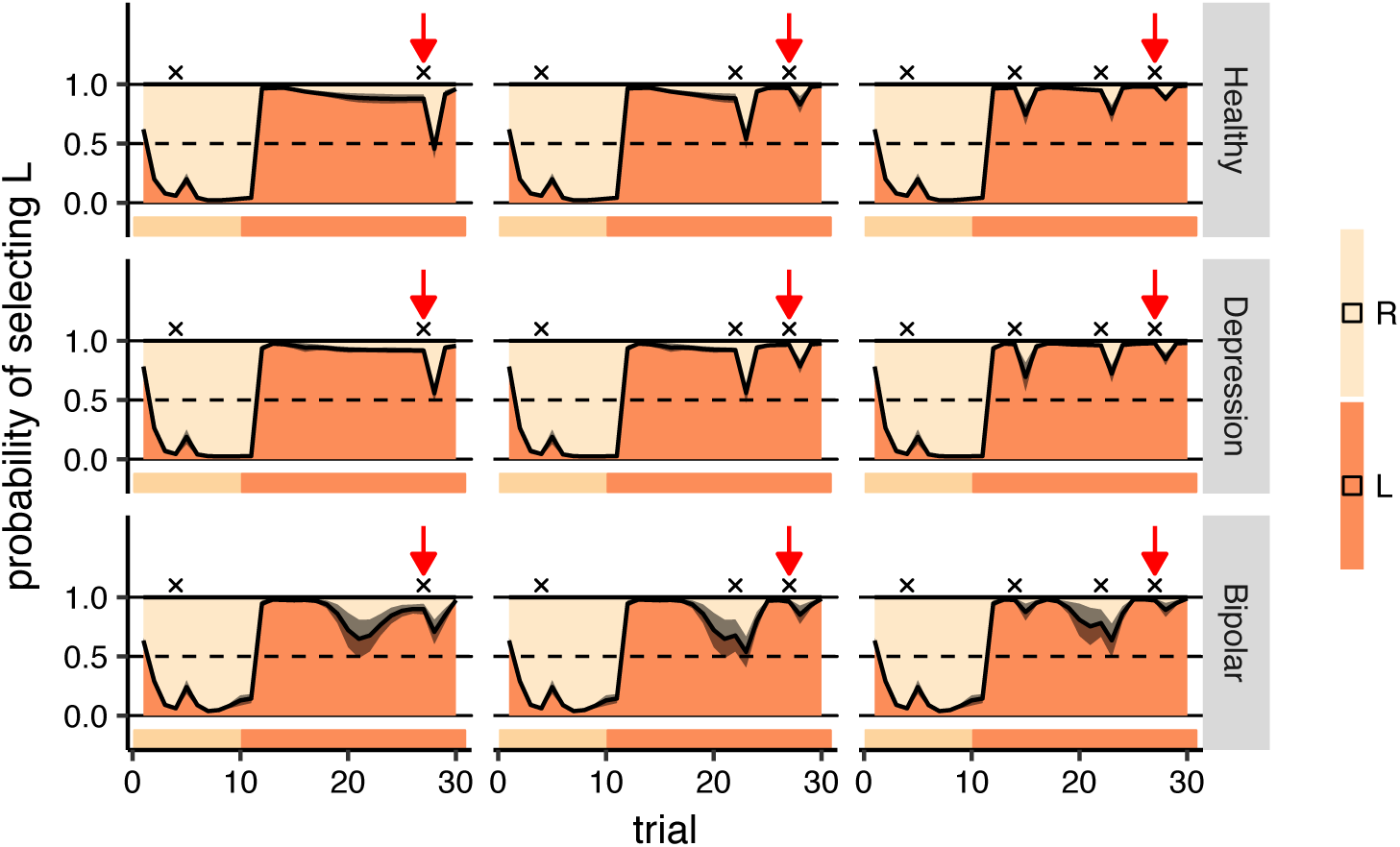
The effect of the initialisation of the network on the off-policy simulations of rnn. The simulation conditions are the same as the ones depicted in Figures 7 and 6. Here, 15 different initial networks were generated and optimised and the policies of the models at each trial were averaged. The gray ribbon around the policy shows the standard deviation of the policies. Each panel shows a simulation for 30 trials (horizontal axis), and the vertical axis shows the predictions of each model on each trial. The ribbon below each panel shows the action which was fed to the model on each trial. In the first 10 trials, the action that the model received was R and in the next 20 trials it was L. Rewards are shown by black crosses (x) on the graphs. See text for the interpretation of the graph.

**Figure S10.**
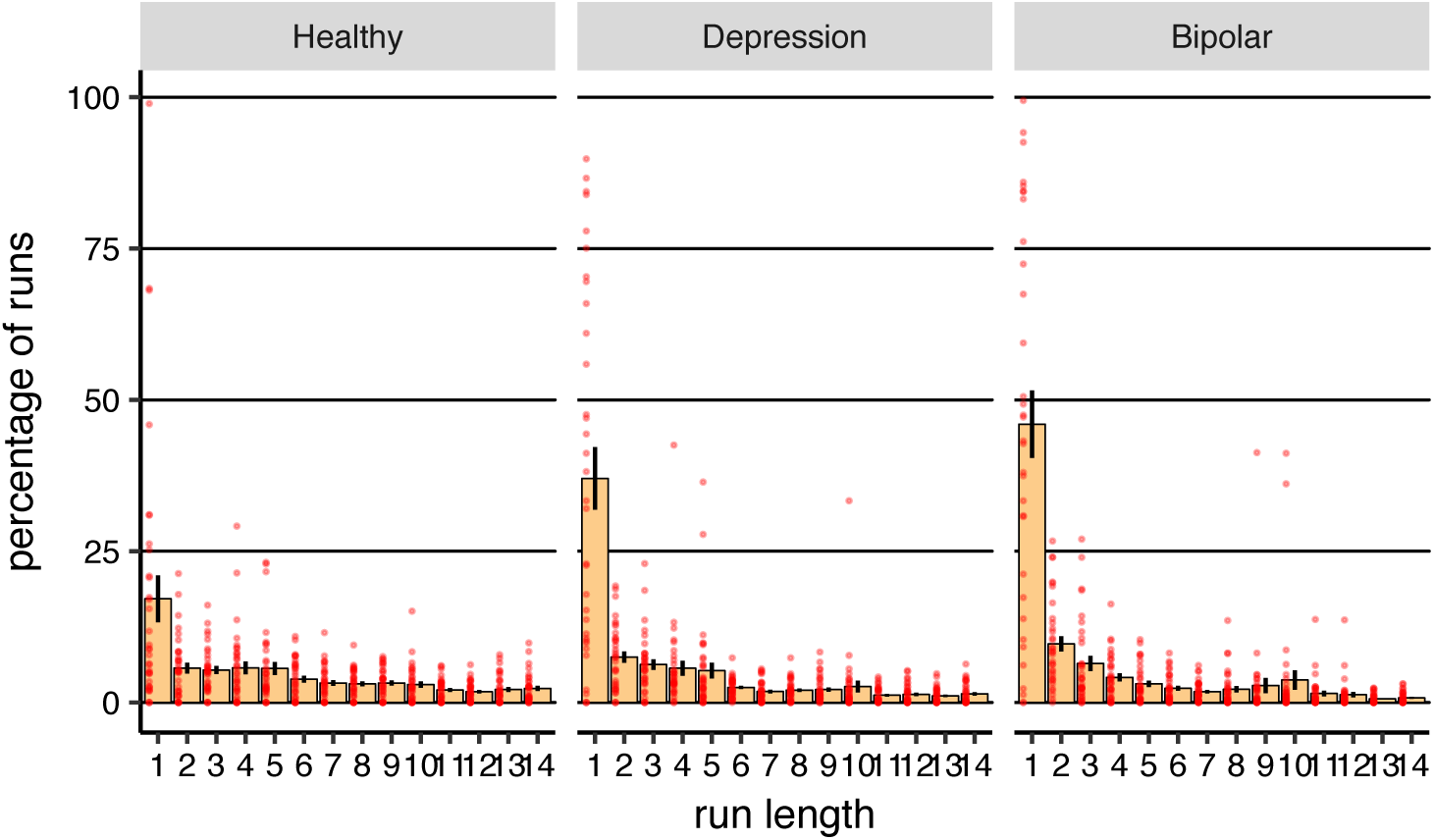
Percentage of each run of actions relative to the total number of runs for each subject. Percentage of each length of run of actions relative to the total number of run of actions in each subject (averaged over subjects). Red dots represent data for each subject, and error-bars represent 1SEM.

**Figure S11.**
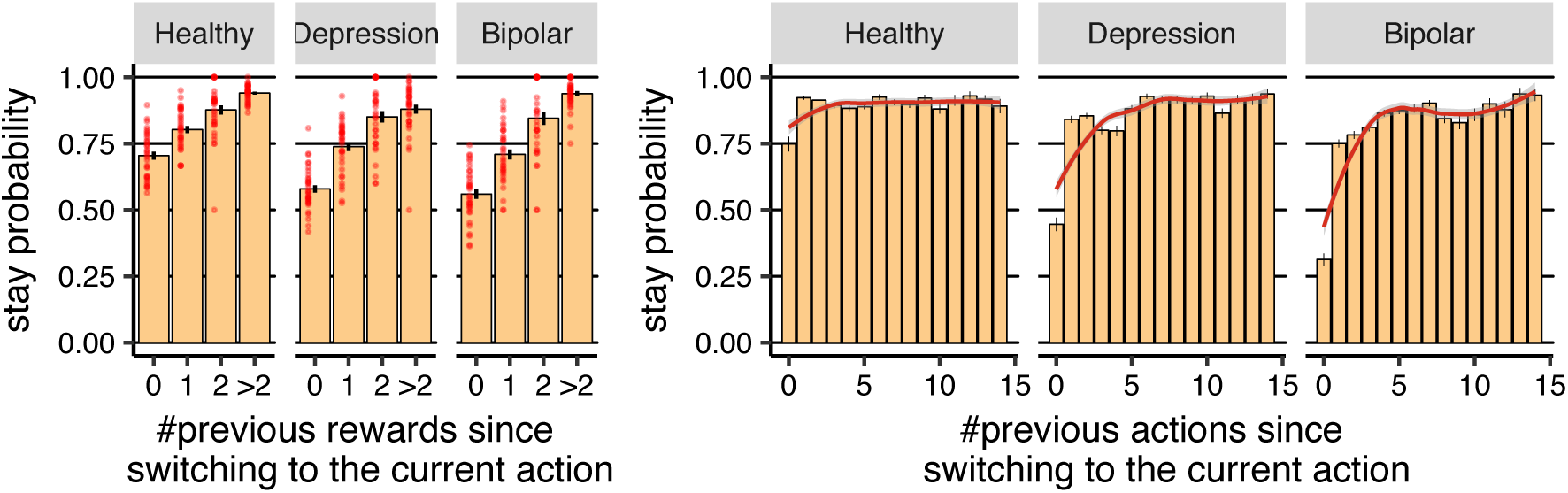
rnn simulations. The graph is similar to Figure 8 but using data from rnn simulations. **(Left-panel)** Probability of staying with an action after earning reward as a function of number of actions taken since switching to the current action (averaged over subjects). Each red dot represents the data for each subject. **(Right-panel)** Probability of staying with an actions as a function of number of actions taken since switching to the current action. The red line was obtained using Loess regression (Local Regression), which is a non-parametric regression approach. The grey area around the red line represents 95% confidence interval. Error-bars represent 1SEM.

**Figure S12.**
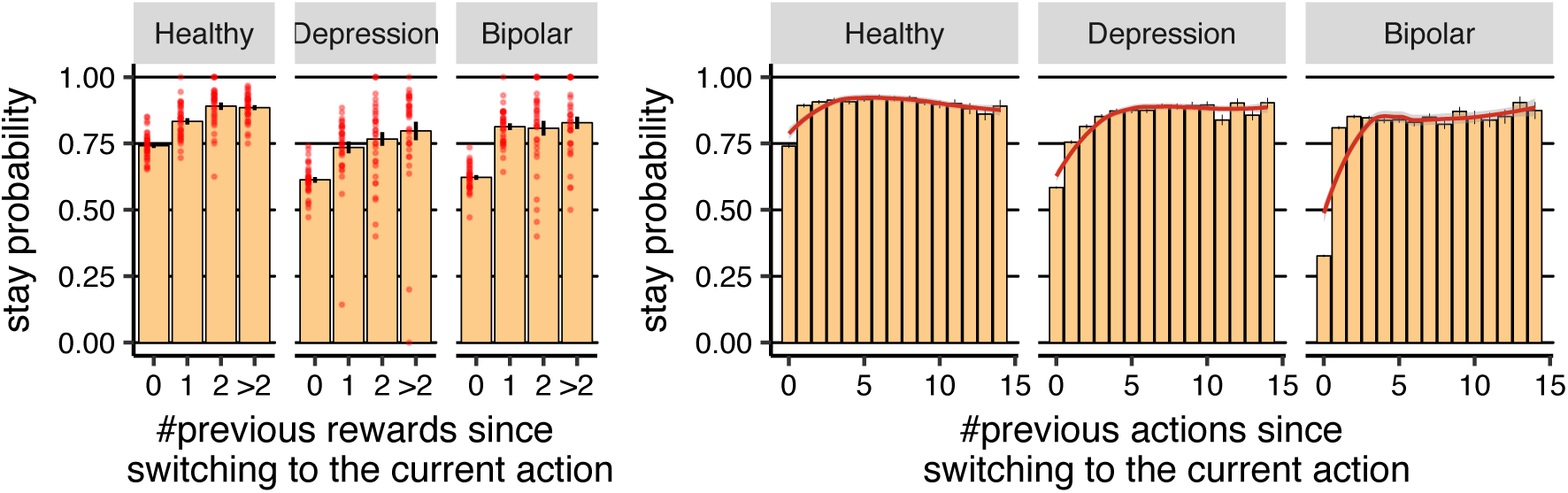
gql simulations (*N* = 2). The graph is similar to Figure 8 but using data from gql simulations with *N* = 2. **(Left-panel)** Probability of staying with an action after earning reward as a function of number of actions taken since switching to the current action (averaged over subjects). Each red dot represents the data for each subject. **(Right-panel)** Probability of staying with an actions as a function of number of actions taken since switching to the current action. The red line was obtained using Loess regression (Local Regression), which is a non-parametric regression approach. The grey area around the red line represents 95% confidence interval. Error-bars represent 1SEM.

**Figure S13.**
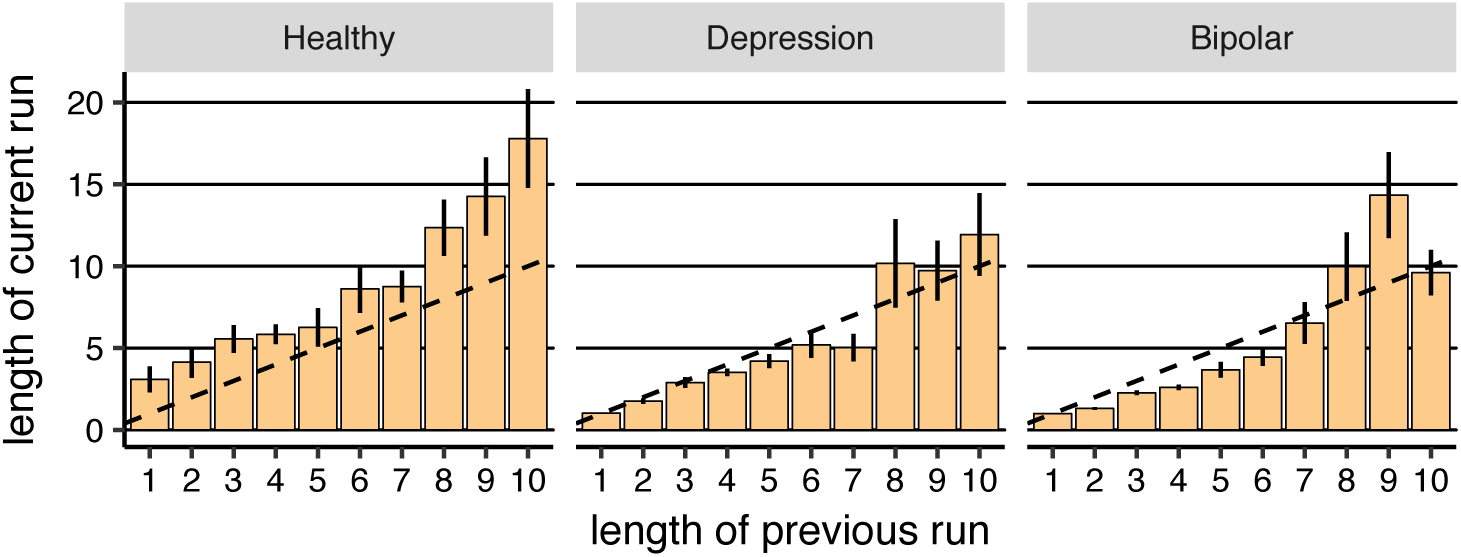
rnn simulations. The graph is similar to Figure 9 but using data from rnn simulations. Median number of actions executed in a row before switching to another action (run of actions) in each subject as a function of length of previous run of actions (averaged over subjects). The dotted line shows the points in which the length of previous and current run are the same. Note that the use of median instead of average was because we aimed to illustrate most common ‘length of current run’, instead of average run length in each subject. Error-bars represent 1SEM.

**Figure S14.**
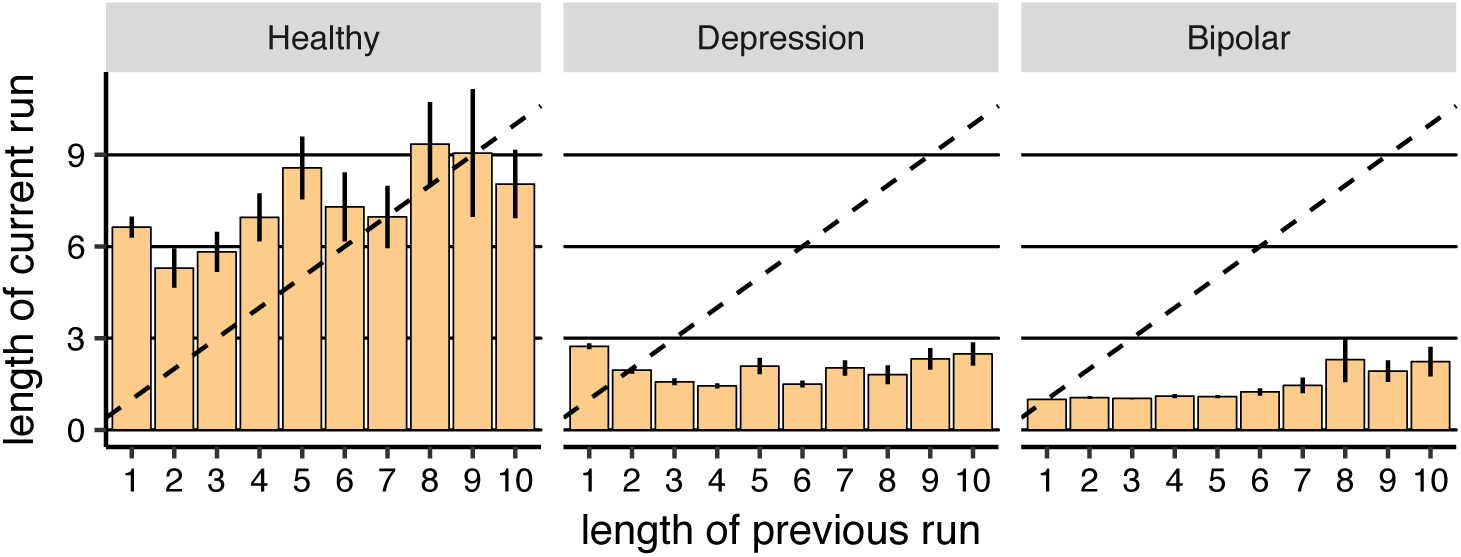
gql simulations (*N* = 2). The graph is similar to Figure 9 but using data from gql simulations with *N* = 2. Median number of actions executed in a row before switching to another action (run of actions) in each subject as a function of length of previous run of actions (averaged over subjects). The dotted line shows the points in which the length of previous and current run are the same. Note that the use of median instead of average was because we aimed to illustrate most common ‘length of current run’, instead of average run length in each subject. Error-bars represent 1SEM.

**Figure S15.**
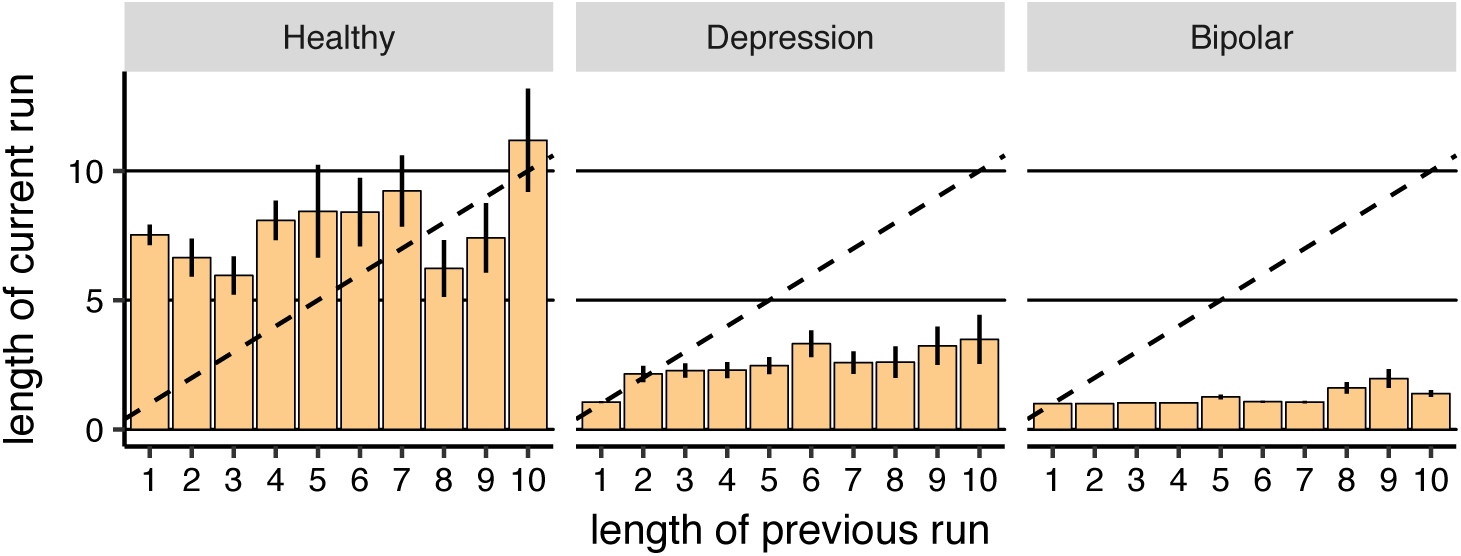
gql simulations (*N* = 10). The graph is similar to Figure 9 but using data from gql simulations with *N* = 10. Median number of actions executed in a row before switching to another action (run of actions) in each subject as a function of length of previous run of actions (averaged over subjects). The dotted line shows the points in which the length of previous and current run are the same. Note that the use of median instead of average was because we aimed to illustrate most common ‘length of current run’, instead of average run length in each subject. Error-bars represent 1SEM.

**Table S1.**
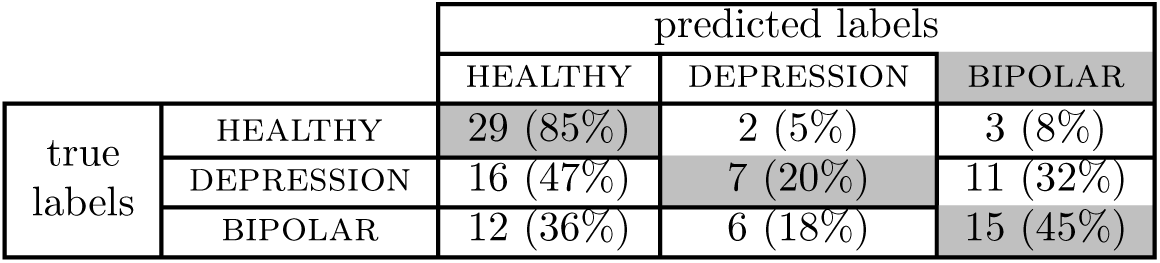
Prediction of diagnostic labels using gql (*N* = 2). Number of subjects for each true-and predicted-labels. The numbers inside parenthesis are the percentage of number subjects relative to the total number of subjects in each diagnostic group.

|  |  | predicted labels |  |  |
| --- | --- | --- | --- | --- |
|  |  | HEALTHY | DEPRESSION | BIPOLAR |
| true labels | HEALTHY | 29 (85%) | 2 (5%) | 3 (8%) |
|  | DEPRESSION | 16 (47%) | 7 (20%) | 11 (32%) |
|  | BIPOLAR | 12 (36%) | 6 (18%) | 15 (45%) |

**Table S2.**
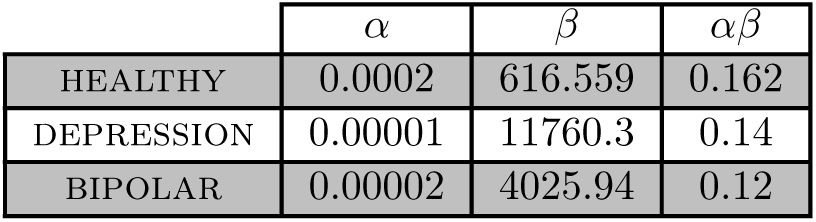
Estimated parameters for ql model.

**Table S3.** Estimated parameters for qlp model.

| | $\alpha$ | $\beta$ | $\kappa$ | $\alpha\beta$ |
| --- | --- | --- | --- | --- |
| HEALTHY | 0.0008 | 98.8895 | 2.0634 | 0.079 |
| DEPRESSION | 0.00003 | 2479.31 | 1.223 | 0.09 |
| BIPOLAR | 0.00008 | 1113.97 | 0.680 | 0.09 |

**Table S4.**
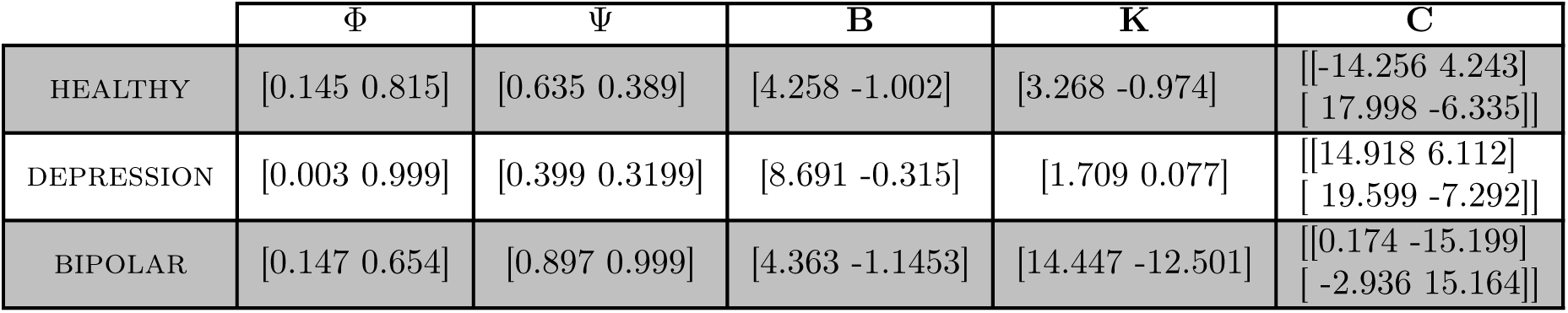
Estimated parameters for gql model with *N* = 2.

**Table S5.** Negative log-likelihood for each model optimized over all the subjects in each group.

|  | RNN | GQL | QLP | QL |
| --- | --- | --- | --- | --- |
| HEALTHY | 9421.6660 | 12939.40 | 14557.44 | 27616.79 |
| DEPRESSION | 13158.1074 | 19735.61 | 23378.65 | 29862.19 |
| BIPOLAR | 12891.3496 | 19363.08 | 24859.15 | 26843.88 |

**Table S6.** Negative log-likelihood for each model. For rnn a single model was fitted to the whole group using ML estimation. For baseline methods (gql, qlp, and ql), a separate model was fitted to each subject, and the reported number is the some of negative log-likelihoods over the whole group.

|  | RNN | GQL | QLP | QL |
| --- | --- | --- | --- | --- |
| HEALTHY | 9421.6660 | 9482.846 | 11529.58 | 26080.13 |
| DEPRESSION | 13158.1074 | 14668.763 | 17837.02 | 28448.94 |
| BIPOLAR | 12891.3496 | 14157.206 | 16912.37 | 25874.56 |

**Table S7.**
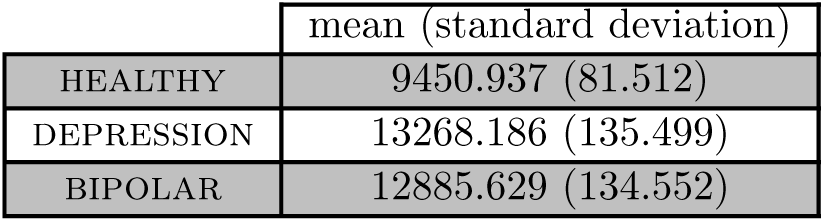
Mean and standard deviation of negative log-likelihood for rnn over 15 different initialisations of the model and optimised over all the subjects in each group.

|  | mean (standard deviation) |
| --- | --- |
| HEALTHY | 9450.937 (81.512) |
| DEPRESSION | 13268.186 (135.499) |
| BIPOLAR | 12885.629 (134.552) |

## Footnotes

1 Albeit more commonly the learner is a human facing a series of learning tasks, rather than a computer model trying to copy the human on a single task.

2 The intercept term was the random-effect at the subject level; action (low reward probability=0, high reward probabilities=1) was the fixed-effect; the dependent variable was the probability of selecting the action.

3 The intercept term was the random-effect at the subject level; and action (low reward probability=0, high reward probabilities=1), groups (healthy, depression/bipolar) and their interaction were fixed-effects; the dependent variable was the probability of selecting the action.

4 The intercept was the random-effect at the subject level; whether reward was earned on the previous trial was the fixed-effect.

5 The intercept term was the random-effect at the cross-validation fold level; model (gql = 1, qlp/rnn = 0) was the fixed-effect.

6 The intercept was the random-effect at the subject level; whether zero rewards or more than two rewards were earned previously was fixed-effect.

7 It can be seen in Figure 3 that the probability of staying with an action was above 50%, irrespective of whether a reward was earned on the previous trial or not. This does not, however, provide evidence for perseveration because the trials were not statistically independent. For example, in late training trials a subject might have discovered which action returns more reward on average and, therefore, stayed with that action irrespective of reward and so without necessarily relying on perseveration.

8 To be consistent with off-policy simulations, only trials on which (i) subjects did not earn a reward on that trial, and (ii) subjects did not earn a reward since switching to the current action, were included in the graph.

9 The intercept was the random-effect at the subject level; the number of times that an action was repeated since switching to the action was the fixed-effect (between zero to 15 times). The dependent variable was the probability of staying with an action.

10 For example, if the executed actions were L, R, R, L, then the length of the first run was 1 (L), the length of the second run was 2 (R, R), and the length of the third run was 1 (L).

11 Note that in on-policy simulations, actions were typically selected probabilistically according to the probabilities that a model assigned to each action. However, in the on-policy simulations presented in this section, in order to get consistent results across simulations actions were *not* selected probabilistically but were chosen based on which action achieved the highest prediction probability.

12 The intercept term was the random-effect at the group level (healthy or bipolar), and the mode of task completion (in the fMRI setting vs. on a computer) was the fixed-effect; the probability of selecting the better key was the dependent variable; see section Statistical analysis for details.

